# Structural evidence for metal ion catalysis in the ribosome

**DOI:** 10.1101/2025.05.16.654626

**Authors:** Jason Hingey, Boris Rudolfs, Daniel B. Haack, Scott A. Strobel, Frank V. Murphy, Simpson Joseph, Navtej Toor

## Abstract

Ribosomes synthesize proteins with an RNA-only active site across all domains of life, yet the details of the catalytic mechanism have remained elusive despite decades of high-resolution ribosome structures. Here, we provide structural evidence for the involvement of two metal ions in peptide bond formation, drawn from ribosome structures spanning bacteria, archaea, and eukaryotes. These metal ions reside in the peptidyl transferase center, one of them adjacent to a universally conserved stack of three base triples, reminiscent of the catalytic triplex in group I/II introns and the spliceosome, which catalyze pre-mRNA splicing. The second metal ion is positioned to stabilize the oxyanion of the tetrahedral intermediate. Metal ion catalysis thus emerges as a recurring mechanistic theme across the central dogma spanning protein synthesis, RNA splicing, and nucleic acid replication.

## Introduction

The ribosome is essential to cellular life, carrying out the intricate task of translating genetic information into functional proteins through peptide bond formation. This process is at the core of the central dogma of molecular biology, in which information encoded in DNA is transcribed to RNA and then translated into proteins that perform nearly all cellular functions. The incredible conservation of the structure and function of the ribosome across all domains of life highlights its evolutionary importance. The first experimental evidence for the RNA-based catalytic function of the ribosome was based on extensive protein removal from ribosomes through proteinase K and SDS treatment while still retaining catalytic activity (*1*). In contrast, activity was abolished by ribonuclease treatment (*1*). This compelling biochemical evidence was later confirmed by high-resolution crystal structures revealing that its catalytic core, known as the peptidyl transferase center (PTC), is remarkably RNA-only, with no protein components directly involved in peptide bond formation (*2*). These findings collectively established that the ribosome functions as a ribozyme, an RNA-based catalyst. Given the essential role of the ribosome in sustaining life, understanding the precise catalytic mechanism of the PTC has been a long-standing goal in molecular biology.

The structural elucidation of the ribosome has been one of the great achievements in biology, yielding atomic-resolution insights into its architecture through x-ray crystallography and, more recently, cryo-electron microscopy (cryo-EM). Over the past two decades, numerous high-resolution ribosome structures from various species have revealed detailed views of its active site, RNA folding, and the positioning of tRNAs and mRNA during translation (*2–6*). These structures have illustrated substrate binding and provided snapshots of the catalytic steps, yet we still lack a clear understanding of how the PTC catalyzes peptide bond formation with such efficiency. This has led to various hypotheses suggesting that the ribosome may utilize alternative strategies, such as substrate positioning (entropic catalysis) (*7*), transition state stabilization through RNA interactions (*8*), contributions from acid-base catalysis by RNA (*9, 10*), or involvement of solvent molecules in proton transfer (*11, 12*). However, there is no clear direct structural evidence to clarify how the chemical environment of the active site might stabilize transition states or lower activation barriers during peptide bond formation.

This gap in understanding is significant, as RNA-based catalysts generally exhibit lower catalytic efficiencies compared to protein enzymes (*13*), prompting questions about whether the ribosome might employ additional strategies to enhance its catalytic power. For example, while RNA alone can catalyze certain reactions, it often relies on metal ions for structural stability and catalytic enhancement. Metal ions are known to play crucial roles in other ribozymes, such as the group I (*14, 15*) and group II introns (*16, 17*), where they are essential for facilitating reactions by stabilizing transition states, activating nucleophiles, and providing electrostatic shielding (*18*). However, until now, there was no structural evidence for the existence of a similar metal-ion-based mechanism that might play a role in peptide bond formation within the ribosome.

In this study, we determined the cryo-EM structure of the cytosolic 80S ribosome from *Chlamydomonas reinhardtii* (*C. reinhardtii*) at a global resolution of 2.7 Å, which indicated the presence of a metal ion within the PTC. We broadened our analysis to include published ribosome structures from many different species and found conservation of metal binding in the PTC across all domains of life. During this process, we identified a second metal ion in the PTC of published ribosome structures containing tetrahedral intermediate analogs. Through the examination of 13 structures from a range of archaeal, bacterial, and eukaryotic species, we identified two metal ions within the ribosomal active site. The first of these metal ions (M1) is positioned alongside universally conserved rRNA residues, in close association with a distinctive stack of three base triples. The second metal ion (M2) is likewise coordinated by universally conserved rRNA residues, but is on the opposite side of the substrate relative to the first metal ion, in a *trans* configuration. Here, we propose that the ribosome uses a novel *trans* two-metal-ion mechanism of catalysis to achieve peptidyl transfer reactions. This mechanism exhibits important parallels to the canonical *cis* two-metal-ion mechanism used by other large ribozymes to carry out phosphoryl transfer reactions.

## Results

### Presence of a metal ion in the peptidyl transferase center

To gain mechanistic insight into the PTC, we determined the cryo-EM structure of the cytosolic 80S ribosome from *Chlamydomonas reinhardtii*. We purified native ribosomes to preserve authentic catalytic states, and collected 4,032 movies on a Titan Krios. The resulting 3D reconstruction achieved an overall resolution of 2.7 Å (**Fig. 1A**). Focusing on the PTC, we looked for any electron density that might hint at reaction intermediates or catalytic cofactors.

**Fig. 1.**
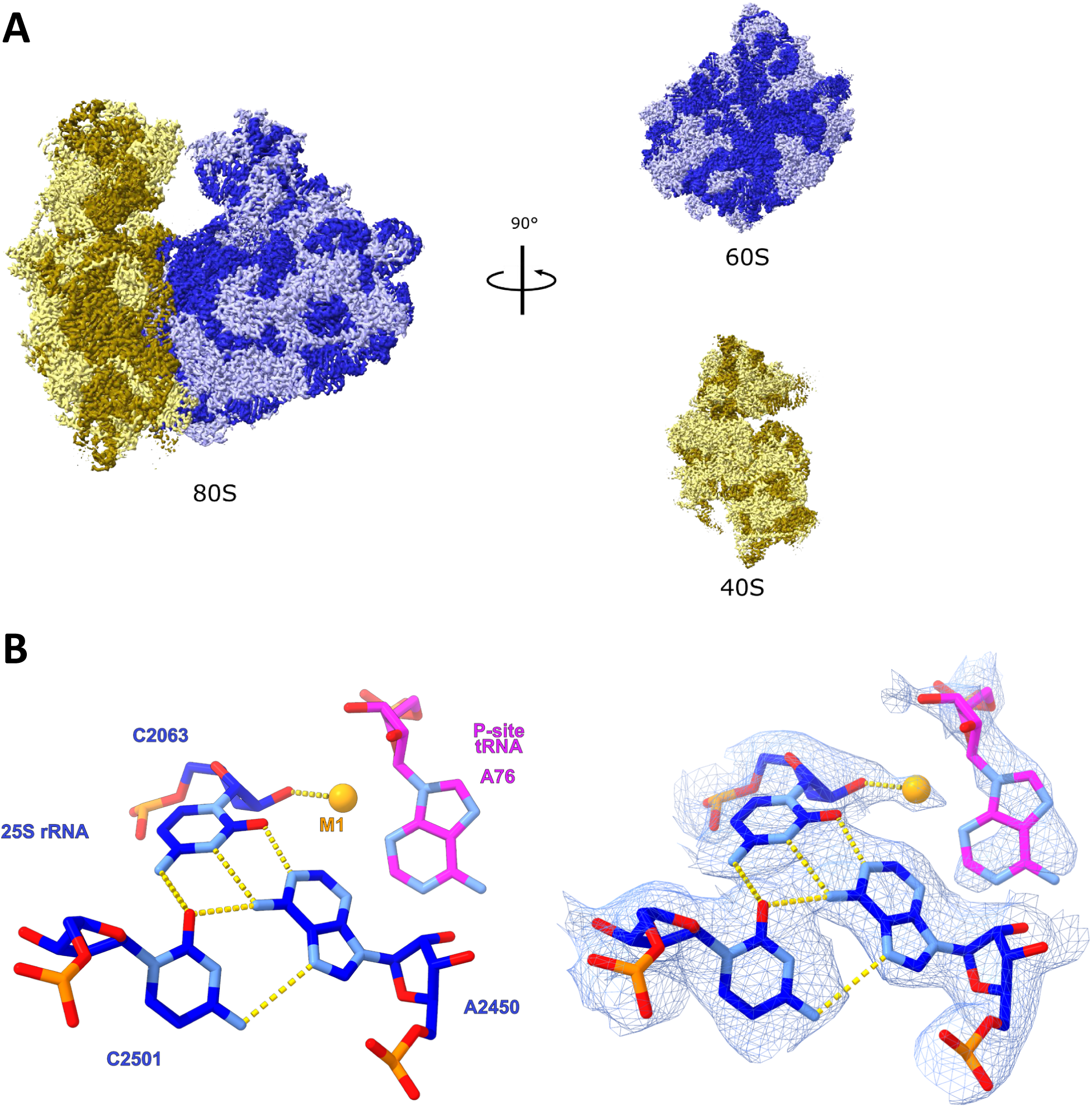
Cryo-EM structure of *Chlamydomonas reinhardtii* cytosolic ribosome. (**A**) Overview of Cryo-EM map of the *C. reinhardtii* 80S ribosome from cryo-electron microscopy. Ribosomal RNA is shown in dark blue and dark yellow. Ribosomal proteins are shown in light blue and light yellow. (**B**) Conspicuous extra density visible in the peptidyl transferase center of the algal ribosome, extending from the 2′-OH of C2063 (using conventional *E. coli* numbering). Metal ion M1 is shown modeled into the density as an orange sphere.

Our initial analysis revealed a density peak in the PTC, consistent with a bound metal ion (**Fig. 1B**). This peak sits adjacent to the 2′-OH of C2063 of the large subunit (using *E. coli* numbering for rRNA residues), which participates in a universally conserved base triple with C2501 and A2450. The non-canonical wobble pair between C2063 and A2450 has been suggested to be protonated at the N1 of A2450 to allow hydrogen bonding to the O2 of C2063(*19*), and is supported by connective density between these two groups in cryo-EM (**Fig. 1B**). The density peak of the metal ion lies in close proximity to the 2′-OH of A76 on the 3′ end of the peptidyl tRNA and the N3 atom of A2451, a universally conserved nucleotide long studied for its hypothesized role in catalysis (*10, 20*) (**Fig. S1**). Given its strategic positioning at the interface between P-site tRNA A76 (bearing the nascent peptide through an O3′ ester linkage) and rRNA residue C2063, we hypothesized that this density possibly corresponds to a metal ion serving as a direct catalytic cofactor in peptide bond formation. We then surveyed ribosome structures spanning ∼25 years to see if this putative metal site was a recurring, conserved feature of the active site.

### Identification of metal ions in x-ray and cryo-EM maps

Metal ion identification in electron density maps becomes reliably achievable at resolutions of 3 Å or better. At this resolution threshold, metal ions exhibit sufficient electron density contrast to enable unambiguous identification and clear discrimination from surrounding RNA backbone density. Lower-resolution maps lack the requisite detail to resolve these individual components, resulting in merged electron density that precludes accurate metal ion localization. Metal ions are identified in crystal structures through an analysis of the coordination distance, coordination geometry, and peak strength. The biophysical properties of inner-shell coordinated metal ions have been previously compiled from an analysis of crystal structures with a resolution better than 1.6 Å (*21*). At this high resolution, the inner-shell coordination distance for magnesium to oxygen ranges from 2.1 to 2.3 Å, and the distance is increased by more than 0.1 Å when magnesium is coordinated to nitrogen (*22*). In structures that exhibit resolution poorer than 2.4 Å, the standard deviation for metal ion distances increases, potentially leading to a longer measured distance from the electron density map (*21*). For example, the standard deviation increases from 0.2 to 0.5 Å as the resolution decreases from 1.6 to 2.2 Å (*21*). Metal ion identity can also be inferred from coordination distance for potassium, which has an inner-shell distance ranging between 2.7 and 3.2 Å (*21*). Similarly, water molecules can be distinguished from magnesium ions by their coordination distance of 2.8 to 2.9 Å (*23, 24*). Coordination geometry with the surrounding protein/RNA ligands is also indicative of metal ion identity. Magnesium exhibits octahedral coordination geometry, whereas potassium has flexible geometry (*21*). Finally, high peak strength with distinct spherical density in a crystallographic *F*_o_–*F*_c_ omit map indicates the presence of an electron-rich metal ion. A more recent development in the generation of electron density maps allows enhancement of weak signals and reduction of noise in the 2*F*_o_–*F*_c_ map. These feature-enhanced maps (FEM) are generated in the absence of any modeled content in the region of interest, and can greatly improve the interpretability of the density (*25*). FEM is part of the standard Phenix structure refinement package (*26*).

Cryo-EM maps usually require higher resolution to resolve metal ions clearly. In our experience, cryo-EM structures need a local resolution of ∼2.5 Å to visualize density for metal ions. We have found that in contrast to x-ray crystallography, it is relatively rare to see discrete metal ion density at a resolution of 3 Å in a cryo-EM map. Cryo-EM and x-ray crystallography differ fundamentally in their scattering mechanisms (*27*), which impacts their sensitivity to atomic charge. Cryo-EM relies on electron scattering, which is influenced by the electrostatic potential of the sample. This includes contributions from both the positively charged atomic nuclei and the electron cloud, making cryo-EM highly sensitive to the net charge of atoms. As a result, cations scatter strongly, while anions may produce weaker or even negative signals at low resolutions. In contrast, x-ray crystallography is based on photon scattering, which depends solely on electron density. Employing both cryo-EM and x-ray crystallography to identify a metal ion position is the most optimal approach.

Work done by Wang et al. (2018 and 2021) shows that the electrostatic potential maps of cryo-EM contain information beyond that found in x-ray crystallography to facilitate identification of metal ions (*27, 28*). Highly positively charged ions like magnesium exhibit an unusual effect in which the inner-shell coordination between a magnesium ion and the polar N or O-containing ligand produces delocalized electrostatic potential, resulting in tubular density connecting these two atoms (*28*). At short interaction distances (e.g., 2.0–2.2 Å), these peaks often merge with neighboring atom signals at low contour levels. This behavior is a useful indicator of inner-shell coordination and assists in metal ion identification. It arises because the strong positive charge is only partially offset by the partial negative charges of the coordinating atoms. In contrast, phosphate group oxygens in the RNA backbone, which carry full negative charges, more effectively reduce the amplitudes of Mg^2+^ peaks, making these peaks more discrete. This density characteristic is helpful in identifying multivalent cations like Mg^2+^ and is not seen for monovalent ions.

### Observation of conserved PTC metal ion M1 in cryo-EM structures

In our *C. reinhardtii* 80S map, the PTC region reaches the sub-2.7 Å local resolution range, enabling identification of the density described above as a likely Mg^2+^. The cryo-EM density appears as a continuous, tubular peak linking the metal ion (M1) to the O2′ of C2063, consistent with inner-shell coordination (**Fig. 1B**). Such tubular connecting density is a hallmark of bound multivalent ions in cryo-EM maps, reflecting delocalized electrostatic potential between a multivalent cation and its ligands (*27, 28*). The M1 peak is only observed in the presence of a P-site substrate. Our high-resolution map shows density for only the 3′ end of the P-site tRNA, due to movements of the anticodon portion of the tRNA during ratcheting of the small subunit as the ribosome translates. Focused classification confirmed this to be the case, revealing the presence of the full P-site tRNA in a variety of conformations (**Fig. S2**).

Having established criteria for recognizing M1 in the PTC, we next asked whether previous high-resolution ribosome structures showed evidence of a similar feature. We restricted our search to cryo-EM structures with local resolution ∼2.5 Å in the PTC to reliably detect M1. We included crystal structures resolved to high resolution, reasoning that if this metal were functionally important, it might appear in prior data even if not reported. Using our *C. reinhardtii* ribosome structure as a guide, we re-examined a diverse set of ribosome structures from bacteria, archaea, and eukaryotes for evidence of a metal ion in the PTC pocket defined by A76, C2063, and A2451. A substrate docked into the P-site seemed to be a prerequisite for the observation of M1, so we focused our search on ribosome cryo-EM structures containing a P-site substrate. This survey yielded the same PTC metal ion in multiple prior structures (*29–33*). In total, we identified an unmodeled density consistent with a Mg^2+^ ion at this site in five additional cryo-EM reconstructions (**Fig. 2**), encompassing ribosomes from bacteria, plants, and mammals. Notably, none of these deposited structures had modeled a metal ion at this location in their original PDB files. In each case, the position of M1 is directly at the site of catalysis: nestled where the peptide bond is formed, between the ester linkage of the peptidyl-tRNA and universally conserved residues of the large subunit rRNA. The consistent presence of this density across evolutionarily distant species and in structures captured under different conditions underscores that this metal-binding site is a conserved feature of the ribosomal active site.

**Fig. 2.**
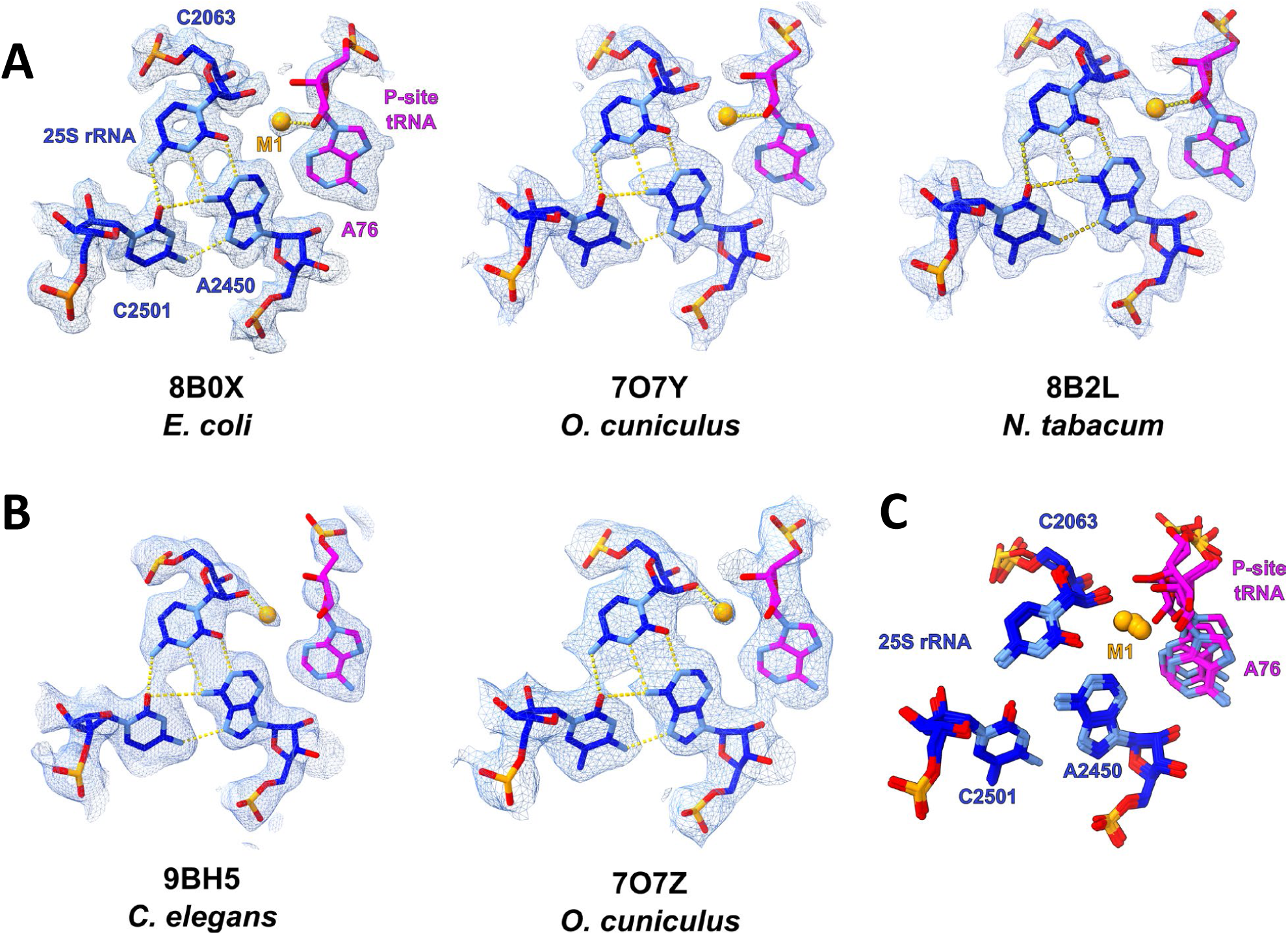
Cryo-EM density for M1 near C2063 and A76. (**A**) Cryo-EM maps showing connecting density between M1 (orange) and the 2′-OH of A76 from the P-site tRNA in three structures. PDB accession codes are indicated. (**B**) Cryo-EM maps showing connecting density between M1 and the 2′-OH of C2063 from the P-site tRNA in two structures. (**C**) An overlay of the PDBs from panels A and B reveals the high degree of conservation in the architecture of this region. M1 moves ∼1 Å between the two states.

Closer inspection of the cryo-EM maps revealed that M1 toggles inner-shell coordination between two nearby 2′-OH ligands. In three of the structures, M1 is positioned closest to the 2′-OH of C2063 at the shortest distance of 2.09 Å, while in three structures M1 is closest to the 2′-OH of A76 from the P-site tRNA with the shortest distance of 2.02 Å (**Table S1**). The metal ion moves ∼1 Å in going from one hydroxyl to the other between these two different sets of structures. In most of these cases, the shorter distance coincides with continuous density connecting M1 to the O2′, indicating inner-shell coordination to the ribose of these universally conserved nucleotides. These two states likely reflect M1 alternating between coordination to A76 or C2063. We infer that M1 can adopt at least two positions in the active site, effectively transferring between the two riboses. Such a shift could correspond to different catalytic states of the ribosome, for instance before and after peptide bond formation (see Discussion). Importantly, in all instances the metal ion is stabilized by at least one inner-shell contact to an RNA 2′-OH group and is in the vicinity of A2451 (**Fig. 3**), which, in addition to A76, is implicated in the hypothesized proton shuttling mechanisms of peptide bond formation (*12, 34*). Based on the ∼2.1 Å coordination distances and continuous density observed in the electrostatic potential maps, we conclude that this density corresponds to Mg^2+^ rather than a monovalent ion or water molecule in each structure. These observations provide structural support for a Mg^2+^ bound in the ribosomal PTC in an analogous manner across species. Furthermore, M1 is only observed in ribosome structures that contain a P-site substrate, which suggests that both hydroxyl groups must be present to form the M1 binding pocket.

**Fig. 3.**
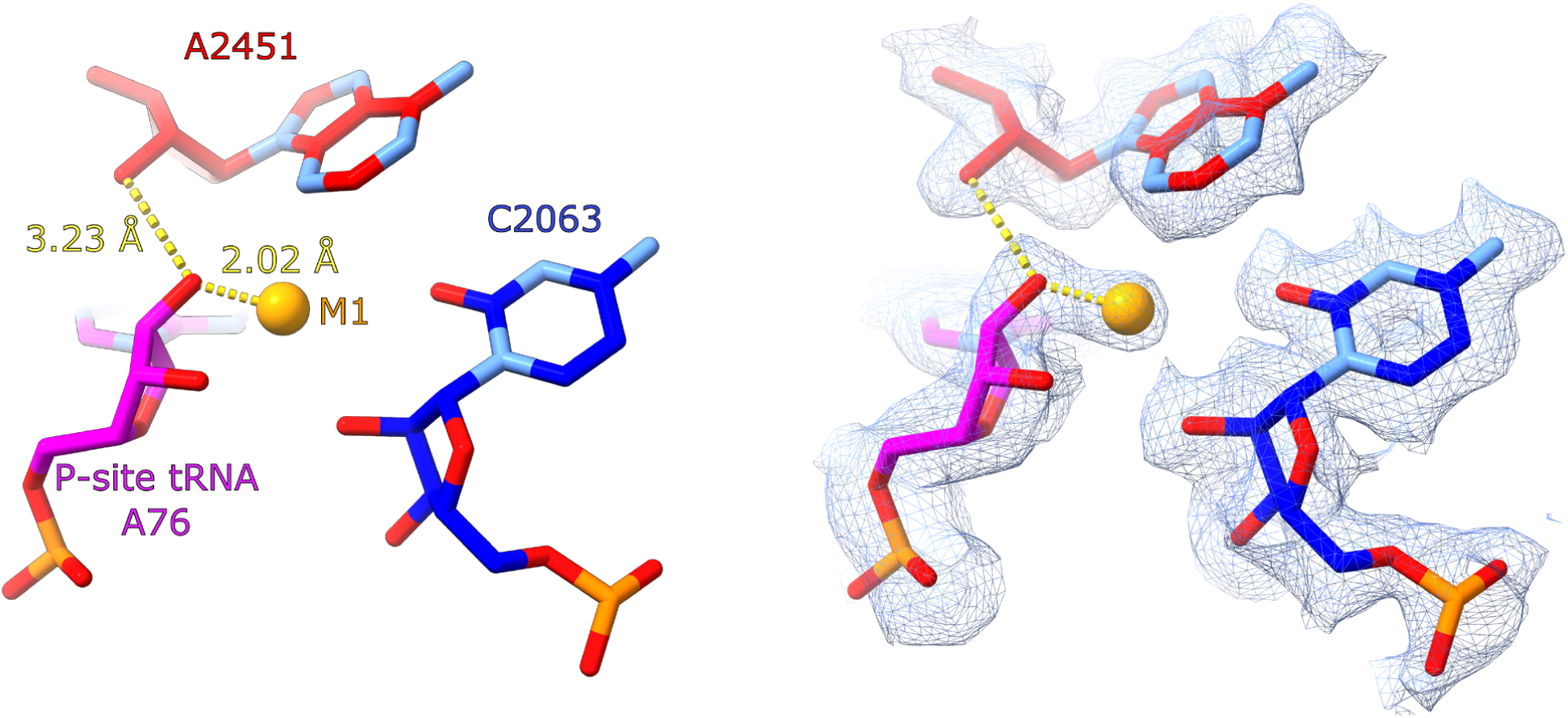
Detailed view of M1 coordinated to A76. The M1 binding pocket in the 1.67 Å cryo-EM map (EMD-15797) associated with PDB deposition 8B0X. The 2′-OH of the P-site A76 inner-shell coordinates M1 (orange) at a distance of 2.02 Å, while also forming a hydrogen bond to the 2′-OH of A2451. In this state, the ribose of A76 exhibits a C2′-endo sugar pucker.

### PTC metal ion M1 in crystal structures of the ribosome

We next turned to x-ray crystal structures of the ribosome to see if M1 is also evident in crystallographic electron density maps. We revisited an early high-resolution structure of the large ribosomal subunit from *Haloarcula marismortui* published in 2002 (*35*). In this 50S crystal structure of *H. marismortui* complexed with a P-site substrate analog (CCA-Phe-caproic acid-biotin) and the antibiotic sparsomycin, we observe strong positive difference density (7.6σ in a *Fo–Fc* omit map) consistent with a metal ion at the site between the P-site substrate and A2451, although nothing was modeled into this region in the original PDB (accession code 1M90) (**Fig. 4**). Additional structures of the *H. marismortui* ribosome were found to have a pair of water molecules modeled into a patch of ambiguous weak density at this site (**Fig. 5A**). We initially tested the use of FEM map generation for PDB 1VQ4 to clarify this signal, which enabled visualization of strong density in close proximity to C2063 (**Fig. 5B**). FEM maps were generated in the absence of a modeled metal ion to avoid bias. The FEM method revealed a spherical peak of electron density located 2.0 to 2.2 Å away from the C2063 2′-OH in these structures (**Fig. 5C, Table S1**). In each of these cases, the Mg^2+^ is coordinated between the 2′-OH of C2063 and the 2′-OH of the P-site tRNA A76 in an arrangement that is consistent with our cryo-EM observations, providing crystallographic evidence that independently corroborates the presence of M1 in the ribosomal active site. The consistency between cryo-EM and x-ray data underscores that this metal ion is not an artifact of one method or a specific sample, but rather an authentic feature of the active site architecture.

**Fig. 4.**
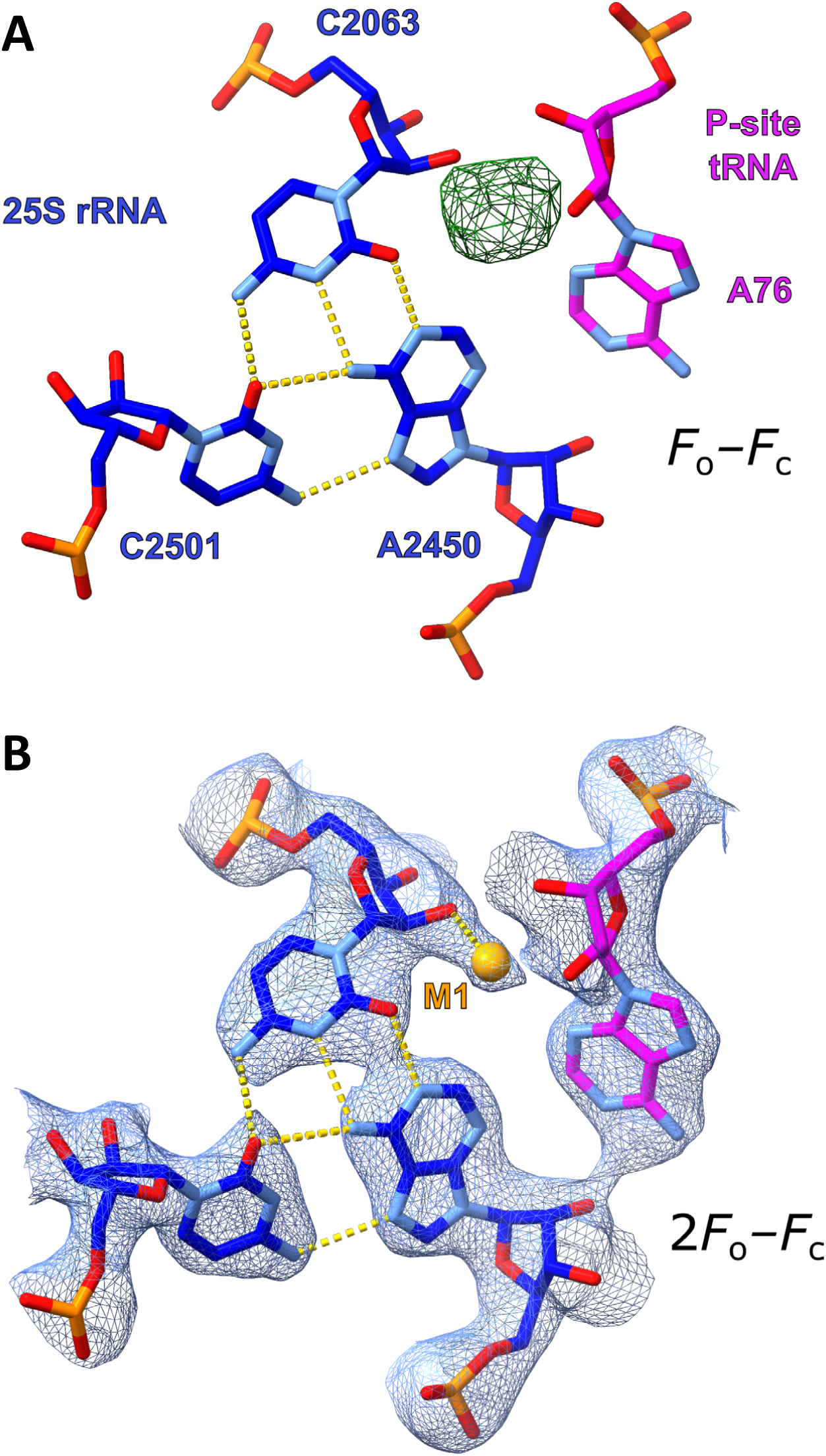
M1 binding pocket in an x-ray crystal structure. (**A**) Crystallographic *F*_o_–*F*_c_ map contoured at 3σ from PDB deposition 1M90, showing strong (7.6σ), spherical positive density consistent with a metal ion. (**B**) Crystallographic 2*F*_o_–*F*_c_ map contoured at 1σ, after refining PDB deposition 1M90 with Mg^2+^ modeled into the electron density.

**Fig. 5.**
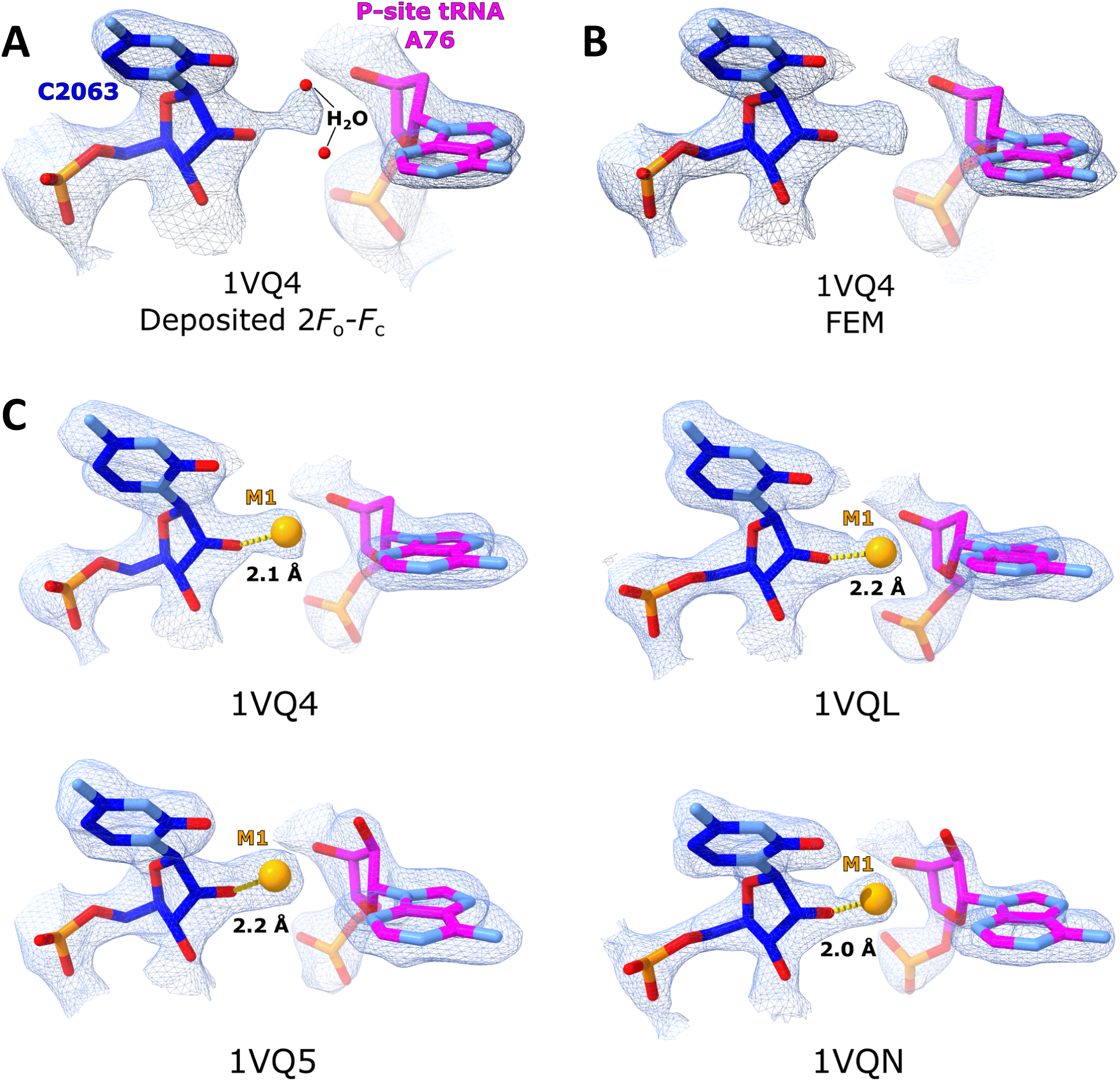
Feature-Enhanced Maps reveal M1 binding pocket in x-ray crystal structures. (**A**) The M1 binding pocket as originally modeled in PDB 1VQ4. Two water molecules are modeled into the region (red spheres) near an ambiguous blip of density between them. The map is contoured at 1.4σ. (**B**) The empty metal ion binding pocket from PDB 1VQ4 after generation of a Feature-Enhanced Map (FEM). The water molecules were removed from the model and no metal ion was modeled prior to creating the FEM in order to avoid any model bias in this region. The map is contoured at 1.8σ. (**C**) Metal ion coordination observed in Feature-Enhanced Maps from x-ray crystallography structures. In all cases, water molecules were deleted for FEM generation as described for panels (A) and (B). PDB accession codes are indicated. Approximate contour levels are as follows: PDB 1VQ4, 1.8σ; 1VQ5, 1.5σ; 1VQL, 1.4σ; and 1VQN, 2.6σ.

### A second metal ion M2 stabilizes the tetrahedral oxyanion intermediate in the PTC

Ribosome structures containing tetrahedral intermediate analogs have previously been modeled with a water molecule positioned near the tetrahedral center (*36*), coordinated by the 2′-OH of U2584 (in some species, post-transcriptionally modified to 3-methyluridine) and the N1 of A2602. This water was proposed to stabilize the developing oxyanion during peptide bond formation (*36*). While stabilization of oxyanion intermediates by divalent metal ions is well-established in many biological systems (*37, 38*), there are no examples of water alone fulfilling this role (*36*). Furthermore, this water molecule is modeled ∼2.5 Å from the 2′-OH of U2584 in these structures, which is outside of the expected 2.8 to 2.9 Å range for a water molecule (*23, 24*) and indicates incorrect assignment.

To probe this assignment, we reprocessed the relevant ribosome structures using FEM refinement. This revealed a well-defined density peak 2.2 to 2.3 Å from the 2′-OH of U2584 and in the vicinity of the tetrahedral oxyanion (**Fig. 6**). The coordination distance is consistent with that expected for a Mg^2+^ ion. M2 is also in close proximity to the oxyanion position on the tetrahedral intermediate analogs.

**Fig. 6.**
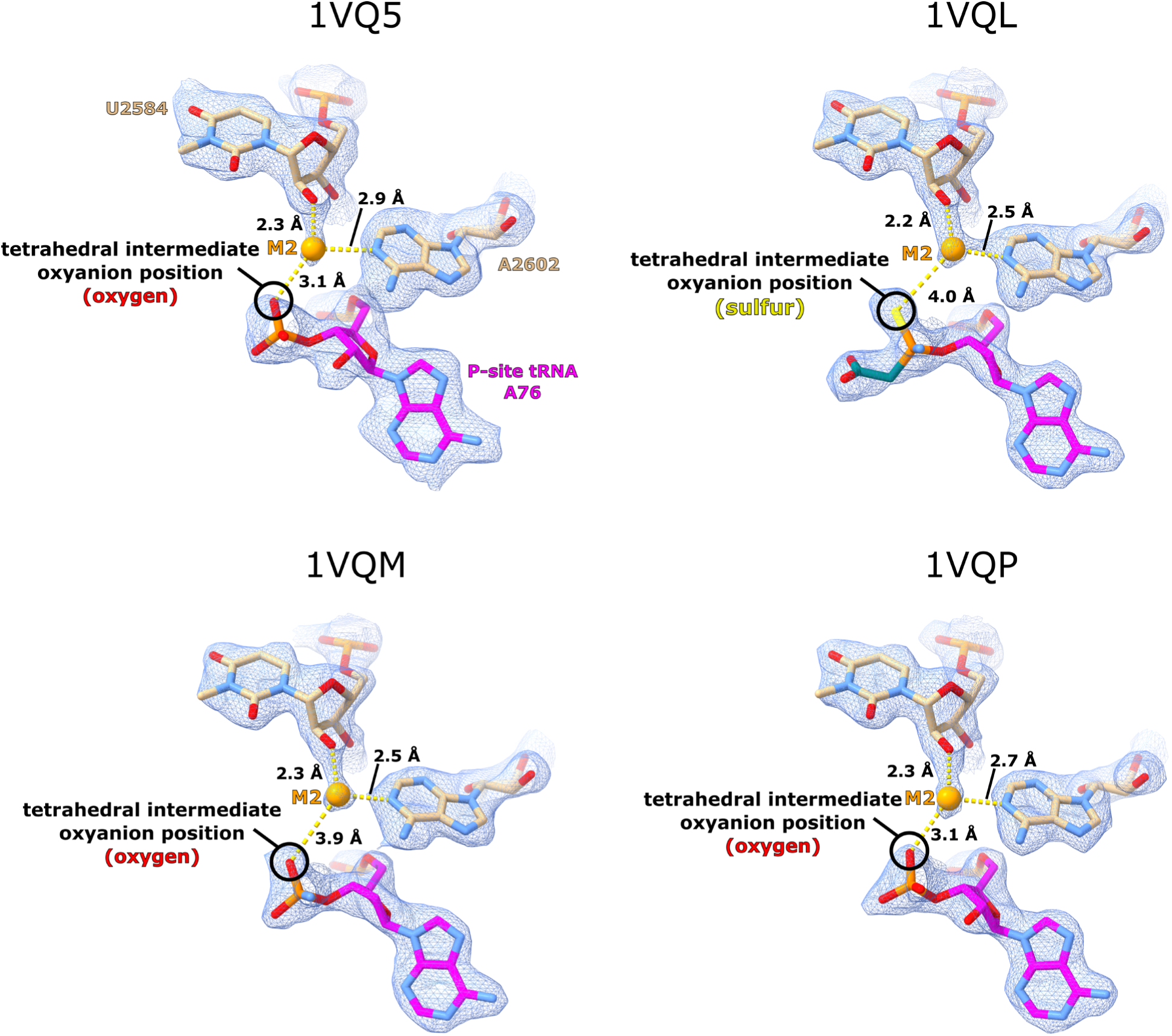
M2 coordination by U2584 and A2602 defines a conserved metal-binding site. M2 is observed coordinating to the 2′-OH of U2584 and the N1 of A2602 in the presence of various tetrahedral intermediate analogs. It lies in close proximity (∼3–4 Å) to the oxyanion position in each of these structures, with the distance impacted by the identity of the atom at the oxyanion position in the analog. The corresponding PDB accession code is indicated above each structure. FEM maps are contoured at approximately 0.9σ, 1.6σ, 1.8σ, and 1.8σ for 1VQ5, 1VQL, 1VQM, and 1VQP, respectively.

### Binding geometry of the PTC metal ions

We observed that there is an adenosine residue in close proximity to each of the two PTC metal ions. When M1 is coordinated to the 2′-OH of A76, the angle formed between the O2′ of A76, M1, and the N3 of A2451 is ∼90° (**Fig. S3A**). In the tetrahedral intermediate analog structures, the angle between the O2′ of U2584, M2, and the N1 of A2602 is likewise ∼90° (**Fig. S3B**). These angles are consistent with the octahedral binding geometry expected for Mg^2+^.

The metal coordination environments in the ribosomal PTC differ fundamentally from those observed in classical two-metal-ion systems. In phosphoryl transfer ribozymes such as group II introns and the spliceosome, both catalytic metal ions are anchored by hard anionic phosphates that provide geometric rigidity and partial charge shielding (*17, 39, 40*). The ribosomal system instead relies on coordination by soft ligands in the form of neutral hydroxyl groups at ribose 2′-OH positions and endocyclic imines of adenosine residues. This involvement of neutral ligands likely preserves greater positive charge on the ribosomal Mg^2+^ ions, enhancing their capacity to stabilize the negative charge buildup during peptide bond formation. The absence of strong anionic ligands explains why these metals appear less localized or variably ordered across structures. Notably, the group I intron represents an intermediate case between these two extremes. While its two catalytic metal ions are stabilized by phosphate coordination similar to group II introns, each metal also forms an inner-shell contact with a 2′-OH group (*15*), establishing a precedent for hydroxyl-mediated metal coordination in ribozyme catalysis. The ribosome extends this strategy further by employing 2′-OH coordination as the primary means of positioning its catalytic metals. In addition, the metal ions in phosphoryl transfer ribozymes are both situated on one side of the scissile phosphodiester bond, in a *cis* configuration. In the ribosome, the two metal ions in the PTC are found on either side of the scissile ester bond, in a *trans* configuration (**Fig. 7**).

**Fig. 7.**
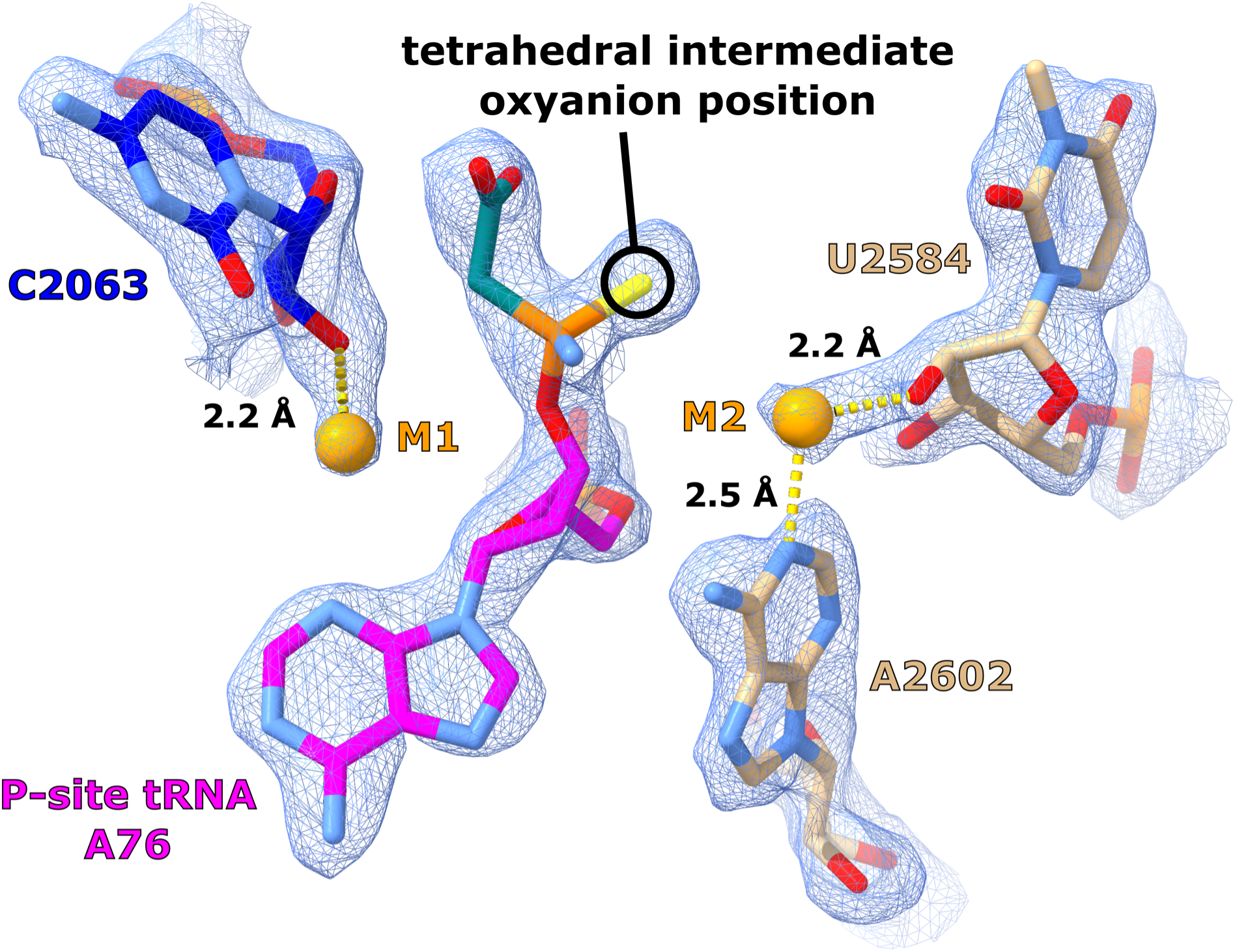
M1 and M2 bracket the site of peptide bond formation in a *trans* configuration. The two catalytic metal ions M1 and M2 are located on opposite sides of the scissile ester bond in a *trans* configuration. In this tetrahedral intermediate analog structure, the oxyanion position is occupied by a sulfur atom (PDB 1VQL, FEM map contoured at 1.4σ).

### Catalytic triplex in the PTC

In analyzing the RNA architecture around the metal ion, we found that the PTC contains a stack of three base triples that is universally conserved. These base triples form a rigid three-dimensional scaffold that forms the binding pocket for M1. This base triple stack is present in all known structures of the ribosome. Such an arrangement of stacked base triples is relatively rare in RNA structures; however, it is common in ribozyme active sites utilizing a two-metal-ion mechanism (*14, 17, 39–41*). Notably, this triple-helix motif exhibits homology to the catalytic triplex found in group I/II introns and the spliceosome (*17, 39–41*). The group I intron is phylogenetically unrelated to the group II intron and the spliceosome, but it also contains three stacked base triples in the active site that form a binding pocket for two catalytic metal ions (*14, 15*). In group II introns and the spliceosome, the triple helix motif in the active site coordinates two Mg^2+^ ions and is dynamic during splicing (*16, 17, 39, 40, 42, 43*). The PTC triples of the ribosome appear to serve an analogous role: they position functional groups (such as the 2′-OH of A76 and the 2′-OH of C2063) in an orientation optimal for binding M1 and for interacting with substrates. This arrangement provides an explanation for the strict conservation of nucleotide identities and the architectural arrangement within the triplex (**Fig. 8A** and **Movie S1**). A side-by-side comparison of the catalytic triplexes found in the ribosome, group II intron, spliceosome, and the group I intron shows structural homology (**Fig. 8**). This parallel suggests an evolutionary convergence on using a triple-helix RNA scaffold to facilitate metal-ion catalysis in ribozymes. In summary, our structural results suggest that the active site of the ribosome is organized to recruit and position two metal ions that may be involved in the catalysis of peptide bond formation, employing structural motifs and strategies strikingly similar to those found in the spliceosome and self-splicing introns.

**Fig. 8.**
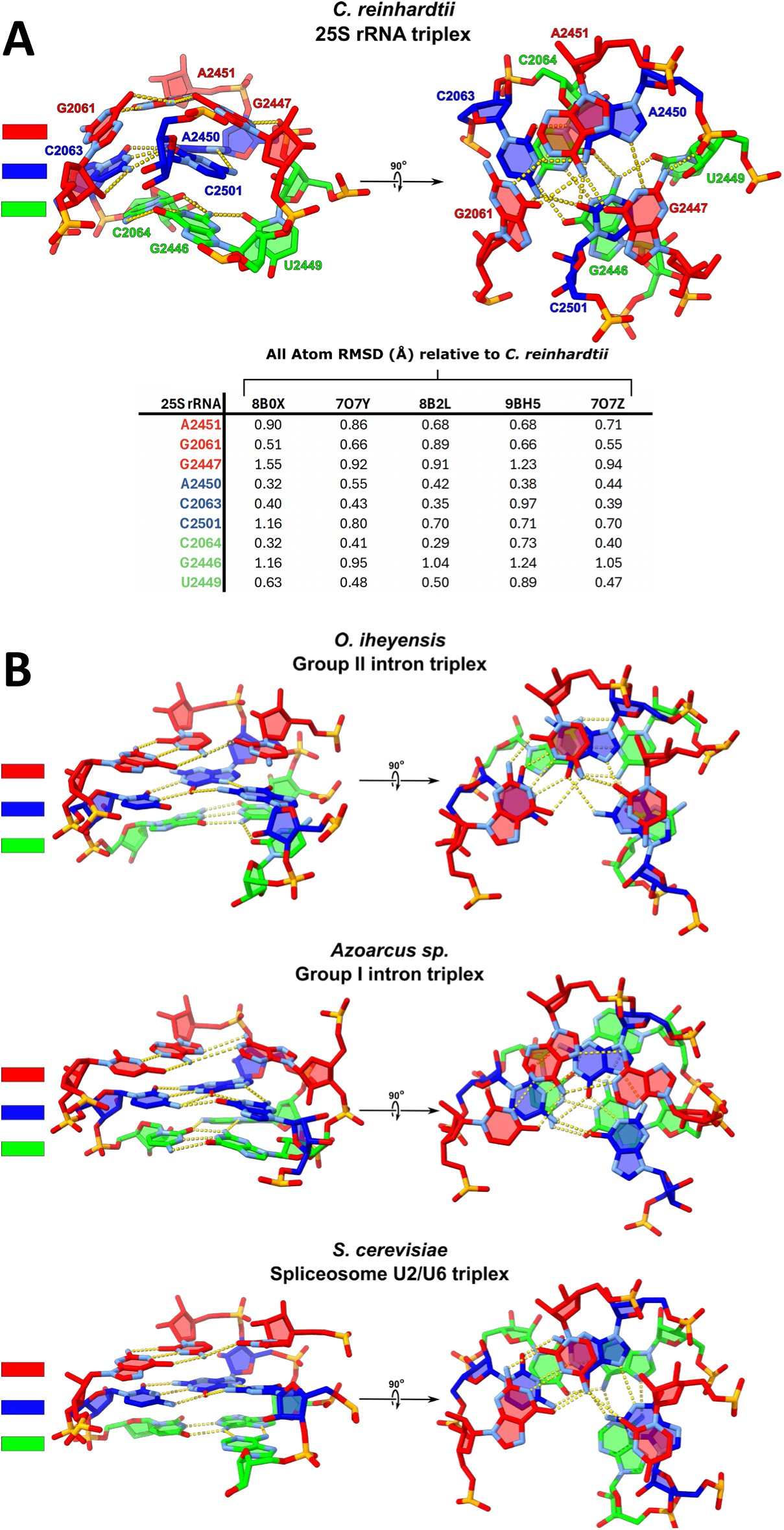
Homologous architecture of the catalytic triplex motif seen in different ribozymes. (**A**) The catalytic triplex of the ribosome as seen in *C. reinhardtii*, with a table of all-atom RMSD values for each nucleotide of the triplex in other cryo-EM structures from bacteria and eukaryotes. Planar base triples are color-coded for clarity. (**B**) Comparison of the catalytic triplex present in group I and group II introns and the spliceosome.

## Discussion

Over two decades of structure-informed research on peptide bond formation in the ribosome have greatly advanced our understanding of its catalytic efficiency, yet a unifying mechanism has remained elusive. Initially, the high-resolution structure of the large ribosomal subunit by Ban et al. (2000) (*2*) spurred numerous hypotheses to explain the strong rate enhancement of the ribosome. Consensus within the field coalesced around the idea of catalysis by approximation, wherein the ribosome functions as a positional catalyst by precisely aligning the A-site and P-site substrates to encourage bond formation (*7*). All enzymes perform this function of substrate positioning in some capacity, as reactive functional groups must be in close proximity to each other for a reaction to occur; however, most enzymes employ additional catalytic strategies for activating functional groups, such as the metal ion catalysis seen in DNA and RNA polymerases (*44–46*). While substrate alignment undoubtedly contributes to the function of the ribosome, this alone has not fully accounted for the observed reaction rate (*34, 47*), especially given that RNA catalysts typically are far less efficient than protein enzymes (*13*). Other proposed mechanisms included general acid-base catalysis by rRNA nucleotides (for instance, the suggestion that the N3 of A2451 could deprotonate the α-amine nucleophile) and electrostatic stabilization of the transition state by the RNA environment (*8, 35*). However, integrating all biochemical and structural data into a single coherent model has proven difficult.

Our structural data revealing two metal ions in the peptidyl transferase center provide a new framework for interpreting decades of biochemical analyses, particularly those concerning nucleotides A2451 and C2063 from the large subunit rRNA and A76 of the peptidyl-tRNA. We observe that M1 directly coordinates to the 2′-OH of the peptidyl-tRNA A76 in multiple high-resolution cryo-EM structures (**Fig. 2A & 3**). This interaction aligns with biochemical studies that identified a critical role for the A76 2′-OH in peptide bond formation, where a 2′-deoxy substitution at this position resulted in up to a 2,000-fold decrease in catalytic rate (*48–50*). In this position, M1 may be involved in activation of the nucleophile, stabilization of the leaving group, or perhaps takes on both of these roles. Inner-shell coordination of M1 to the 2′-OH of A76 lowers the pK_a_ of this functional group, which could facilitate the transfer of a proton to the O3′ position and resolve the tetrahedral intermediate, as hypothesized by Weinger et al. (2004) (*48*). This particular proton transfer event has been widely suggested to produce the 3′-OH leaving group (*36, 51, 52*). Metal-assisted nucleophilic activation could also feasibly occur. Although we have not identified any structure that captures the simultaneous three-way interaction between M1, the 2′-OH of A76, and the α-amine of the incoming amino acid, M1 coordination to the A76 2′-OH is observed in several cryo-EM structures (**Figs. 2A & 3**), while hydrogen bonding between the α-amine and the A76 2′-OH has been suggested on the basis of biochemical experiments and observed in multiple crystal structures (**Fig. S4**) (*34–36, 48, 49, 52, 53*). If both of these interactions can occur simultaneously, the resulting spatial arrangement could allow M1-mediated activation of the α-amine nucleophile.

Notably, A2451 and its interaction with A76 has also been implicated in maintaining the precise geometry of the PTC (*20*). Biochemical evidence has shown that the 2′-OH of A2451 is critical for peptidyl transferase reaction (*20, 54, 55*). Given the apparent hydrogen bonding between the 2′-OH of A76 and A2451 (**Fig. S4**), the 2′-OH of A2451 may be involved in maintaining the ribose sugar pucker conformation of A76, which in turn could influence the capacity of the 2′-OH of A76 to engage in metal ion coordination. This structural network suggests a means by which hydrogen-bonding interactions within the PTC couple active site geometry to metal ion coordination.

The variable position and connective density exhibited by M1 with both the 2′-OH of A76 and the 2′-OH of C2063 (**Fig. 2**) suggest that this metal ion may sample distinct coordination environments during different stages of the elongation cycle. The alternative coordination of M1 to the 2′-OH of C2063 that we observe in both cryo-EM (**Fig. 2B**) and x-ray crystallography (**Fig. 4 & 5**) may represent a holding position as the active site resets during translocation. This interpretation is supported by earlier analyses showing that nucleobase mutations or ribose modifications at C2063 result in relatively minor effects on single-turnover chemistry (*56*) but significantly compromise multi-turnover peptide synthesis (*57*). This result suggests that while C2063 is not required for the chemical step of peptide bond formation, it may facilitate subsequent rounds of catalysis, potentially by contributing to metal ion repositioning or retention during translocation. Taken together, these biochemical findings support a model in which M1, coordinated by the A76 2′-OH during catalysis and by C2063 at other stages of the elongation cycle, plays a dynamic role in peptide bond formation.

The presence of a P-site substrate appears to be required for the binding pocket to retain M1; however, analysis of multiple cryo-EM reconstructions containing P-site substrates reveals that not all of these structures display metal ion density. For instance, among the five maps published alongside PDB deposition 8B0X, only one shows clear density for the metal. Comparison of two high-resolution maps (1.55 Å and 1.67 Å) from this deposition reveals the structural basis for this variability. In the 1.55 Å map (EMD-15793), the A76 and C2063 2′-OH groups are hydrogen-bonding with each other at a distance of 2.65 Å, while the hydrogen bond between the A76 and A2451 2′-OH groups is not maintained (**Fig. S5**). The hydrogen bond formed between the 2′-OH groups of A76 and C2063 leaves insufficient space for metal ion coordination. In contrast, the 1.67 Å map (EMD-15797) shows the 2′-OH of A76 hydrogen-bonding to the 2′-OH of A2451. This configuration allows M1 to coordinate to the A76 2′-OH at a distance of 2.02 Å, which is ideal for inner-shell Mg^2+^ coordination (**Fig. 3**). This structural heterogeneity suggests that the metal binding pocket exists in equilibrium between occupied and unoccupied states.

The second metal ion, M2, is detected only in crystal structures that contain a tetrahedral intermediate analog in the PTC (**Fig. 6**). Although these analogs differ in geometry and charge density compared to the native tetrahedral intermediate (notably with the substitution of a phosphorus atom for the carbon that would participate in the peptide bond), the complexes reported by Schmeing et al. (2005) (*36*) remain the highest-resolution structural approximations of the transition state available to date. In these structures, M2 was originally modeled as a water molecule at a distance of only 2.5 Å from the 2′-OH of the universally conserved U2584, which is too close for a water. The authors had proposed that a water molecule at this position stabilizes the oxyanion, despite acknowledging the absence of any documented precedent for water-mediated stabilization of a transition state oxyanion (*36*). We find that the coordination distance and geometry are consistent with that expected for a Mg^2+^ ion. The location of M2 is consistent with a role in electrostatic stabilization of the high-energy, negatively charged tetrahedral species, and such stabilization is an expected and well-established role of a catalytic metal ion (*37, 38*).

Despite the notable parallel of employing two catalytic metal ions, the proposed *trans* two-metal-ion mechanism of peptidyl transfer in the ribosome is distinct from the *cis* two-metal-ion mechanism seen in ribozymes that facilitate phosphoryl transfer reactions such as the group I and group II introns. Rather than both metal ions being situated on one face of a scissile phosphodiester bond (*cis*), the two metal ions that we observe in the PTC are positioned on opposite sides of the scissile ester bond of the P-site tRNA (*trans*) (**Fig. 7**), such that only M1 is coordinated by the catalytic triplex. The *trans* arrangement additionally necessitates a larger distance between the two catalytic metals ions, which lie approximately 8.5 Å apart, compared to the conserved distance of ∼4 Å in *cis* two-metal-ion systems (*37*). The coordination environments of the two metals in the PTC are also different from those observed in *cis* two-metal-ion systems, where Mg^2+^ ions are coordinated by hard ligands such as phosphates or carboxylate groups. In the PTC, both metals are instead coordinated by a hydroxyl group and an endocyclic imine from a nearby adenosine residue (N3 of A2451 for M1, and N1 of A2602 for M2), which are neutral functional groups. The observation that M2 of the PTC is only ordered in structures containing a tetrahedral analog suggests that this site is disordered, becoming structured only when the intermediate forms. Together, these findings are consistent with a model in which the ribosome employs a *trans* variant of the two-metal-ion strategy, with M1 stabilizing the leaving group and M2 stabilizing the tetrahedral oxyanion intermediate.

Remarkably, a Tetrahymena group I intron ribozyme, classically known for catalyzing self-splicing via phosphodiester transfer, has been shown to catalyze a peptidyl ester hydrolysis reaction under artificial conditions (*58*). In this study, the active site of the group I intron was re-engineered to bind an N-formyl-methionyl-tRNA fragment and cleave the aminoacyl ester bond. This reaction is analogous to the reverse of peptide bond formation. They found that catalysis by the ribozyme required Mg^2+^ ions and intact catalytic core elements, showing that RNA can use metals to promote chemistry at a carbon center, not just at a phosphorus. This work greatly expanded the known catalytic repertoire of RNA and hinted that the first aminoacyl-tRNA synthetase could have been an RNA. The observation that an unrelated ribozyme could be adapted to catalyze a reaction on an aminoacyl ester only in the presence of magnesium provides an interesting parallel to our finding that the ribosome uses magnesium to assist in processing its aminoacyl ester substrates. The aminoacyl esterase activity of the group I intron can be seen as functionally reminiscent of the peptidyl transferase activity of the ribosome. This example demonstrates that RNA can catalyze chemistry at an aminoacyl ester using magnesium, thus supporting the plausibility of a similar mechanism in the ribosome.

Zhang and Cech (1997) report the *in vitro* selection of a ribozyme that catalyzes peptide bond formation via the same chemistry of peptide bond formation used in the ribosome (*59*). In this system, an N-blocked methionine linked to adenosine through an ester bond acts as an aminoacyl donor and reacts with phenylalanine tethered to the 5′ end of the ribozyme to yield a Met–Phe dipeptide. The ribozyme enhances the uncatalyzed reaction rate by ∼10^6^-fold but exhibits no detectable activity in the absence of Mg^2+^, indicating a strict requirement for divalent metal ions (*59*). Catalytic rates increase sharply at elevated Mg^2+^ concentrations, reaching maximal activity above 100 mM MgCl_2_ (*59*). The steep magnesium dependence led Zhang and Cech to suggest that Mg^2+^ may play a direct catalytic role rather than solely stabilizing the RNA fold. Chemical probing and secondary structure analysis by Zhang and Cech (1998) identified conserved RNA motifs homologous to the peptidyl transferase loop of 23S rRNA, including at least eight of the nine nucleotides of the triplex and the M2-coordinating U2584 (*60*). The remarkable secondary structure homology and Mg^2+^ dependence observed here suggest that divalent metal ions may be conserved catalytic cofactors in both artificial and biological RNA-based peptide bond formation.

It is also insightful to compare the mechanism of the ribosome with that of purely protein enzymes that catalyze similar reactions. One example is carnosine dipeptidase 2 (CNDP2), a human enzyme that, in addition to being a dipeptidase, can condense small metabolites with amino acids (for instance, forming N-lactoyl-phenylalanine from lactate and phenylalanine) to form a peptide bond linkage (*61*). More recently, it has been found to catalyze the condensation of beta-hydroxybutyrate (BHB) and phenylalanine to form BHB-Phe (*62*). In both cases, an amino group is activated to attack a carboxyl group, leading to the formation of a peptide bond, similar to the peptide bond formation that occurs in the peptidyl transferase center of the ribosome. The active site of CNDP2 contains two Mn^2+^ ions that engage in a two-metal-ion mechanism (*62*), directly paralleling our hypothesis that the ribosome relies on divalent metal ions, for catalysis. Just as the ribosome positions substrates and harnesses Mg^2+^ to lower the activation barrier, CNDP2 uses Mn^2+^ to orient its substrates and stabilize the transition state. Metalloproteases are another class of protein enzymes that use catalytic metal ions to engage in peptide bond chemistry. Metalloproteases have either mononuclear or binuclear metal ion active sites to catalyze the cleavage of peptide bonds. The existence of CNDP2 and metalloproteases demonstrates that metal-ion catalysis can achieve peptide bond formation and cleavage, respectively. These examples indicate that metal ion-mediated peptide chemistry is a fundamental biochemical strategy found in nature.

The classic two-metal-ion mechanism, first articulated for protein enzymes and ribozymes by Steitz and Steitz (1993) (*37*), is a recurring strategy in biology (*15, 17, 39, 40, 44–46*). This mechanism contributes to precise substrate positioning and transition-state stabilization in many enzymes involved in the processing or synthesis of nucleic acids. In DNA/RNA polymerases, one Mg^2+^ coordinates the 3′-OH of the primer and deprotonates it to generate a strong nucleophile, while the second Mg^2+^ coordinates phosphate of the incoming nucleotide and stabilizes the developing negative charge as the transition state forms (*44–46*). This two-metal-ion mechanism also exists in group I/II introns and the spliceosome, which catalyze phosphoryl transfer: one metal ion activates the attacking 2′ or 3′-OH, and another stabilizes the leaving group and transition state (*15, 17, 37, 39, 40*). We have identified two metal ions in the PTC. Coordination distances and binding geometry indicate that both metal ions are Mg^2+^ ions. We propose that M1 may either stabilize the leaving group, assist in activating the α-amino nucleophile to attack the ester of the nascent peptide, or perhaps perform both of these functions, while M2 stabilizes the oxyanion in the transition state. By demonstrating the structural presence and likely catalytic function of two metal ions in the peptidyl transferase center active site of the ribosome, we hope to encourage researchers to revisit classical models of ribosome function and to design experiments that directly probe the role of these metal ions.

Our findings not only provide new insights into ribosomal catalysis but also suggest a broader evolutionary conservation of metal-ion-based mechanisms across the central dogma of molecular biology. Our study provides a new lens through which to view the ribosome: not merely as a passive scaffold for bringing reactants together, but as a ribozyme that coordinates catalytic metal ions to achieve proficient peptide bond synthesis. This perspective unifies the mechanism of peptidyl transferase in the ribosome with those of polymerases, the spliceosome, and self-splicing ribozymes, suggesting that nature has repeatedly solved similar chemical challenges using the same fundamental toolkit of divalent metal ions. Ultimately, this understanding of ribosomal chemistry could shed light on how the translational apparatus evolved from simpler RNA-only precursors. The discovery of metal ions in the ribosome PTC thus bridges a long-standing gap in molecular biology, connecting the mechanistic dots from the RNA World to modern protein synthesis.

A significant limitation of this study lies in the inherent difficulty of biochemically validating metal ion coordination by the 2′-OH groups through conventional chemical substitution approaches. Unlike the phosphate oxygen substitutions used in phosphorothioate interference experiments that successfully identified catalytic metal binding sites in the group I (*63*) and group II introns (*64–66*) and spliceosome (*67*), modifications to the ribosomal 2′-OH positions present unique challenges due to their dual functionality, especially in the case of A76. The 2′-OH of A76 is implicated in metal coordination and plays an essential catalytic role in the proton shuttling mechanism during peptide bond formation. This makes it difficult to decouple metal-binding function from its chemical participation in catalysis through simple substitution strategies. While replacing a 2′-OH with a 2′-SH group might maintain coordination with thiophilic metal ions such as Mn^2+^, such modifications would fundamentally alter the proton transfer dynamics and potentially disrupt the precise hydrogen bonding network that maintains the C2′-endo conformation of the A76 ribose. In addition, thiol substitution would preclude the hypothesized interaction between this 2′ position and the α-amino group for nucleophilic activation. The technical challenges of introducing specific functional group substitutions within the large subunit ribosomal RNA through either chemical synthesis or enzymatic incorporation also severely limit the experimental approaches available for direct biochemical validation. These constraints necessitate reliance on the convergent structural evidence presented here from multiple high-resolution structures across diverse organisms, though future development of novel chemical biology approaches may eventually enable more direct functional interrogation of this proposed metal binding site. We hypothesize that the conserved catalytic triplex observed in the ribosome, spliceosome, and group I/II ribozymes represents a primordial RNA enzyme motif dating back to the RNA World. This triple-helix tertiary interaction recurs at the active sites of all these molecular machines, where it positions reacting groups and coordinates catalytic metal ions. In group II introns (and similarly in the spliceosomal U2/U6 snRNA core), an AGC base-triad within this triplex binds two Mg^2+^ ions to facilitate a two-metal-ion mechanism for phosphoryl transfer (*17, 39, 40*). Likewise, a base triple sandwich stabilizes the catalytic core of a group I ribozyme (*14, 15*). Even the peptidyl transferase center of the ribosome is buttressed by analogous base triples and metal ion coordination in the absence of proteins. We propose that this simple, yet versatile motif may have constituted the core of an ancestral replicase that could align substrates and activate them through coordinated Mg^2+^ chemistry. This mechanism could have enabled template-directed RNA polymerization, thereby initiating the self-perpetuating genetic cycles necessary for the emergence of the first life forms in the RNA World. It is intriguing to speculate that the catalytic triplex motif found in the heart of the ribosome, spliceosome, and group I/II intron may be the primordial engine that started life on earth.

## Acknowledgments

We would like to thank the Cal-Cryo Facility at UC Berkeley and Daniel Toso for assistance with data collection. We thank David Hiller for comments and discussion regarding the manuscript.

## Author contributions

Conceptualization: JH, DBH, SAS, SJ, NT

Funding acquisition: FVM, SJ, NT

Investigation: JH, NT

Methodology: JH, BR, DBH, FVM, NT

Supervision: SJ, NT

Visualization: JH, BR, DBH, SAS, NT

Writing – original draft: JH, NT

Writing – review & editing: JH, BR, DBH, SAS, FVM, SJ, NT

## Data and materials availability

The cryo-EM maps and associated atomic coordinate model for the *Chlamydomonas reinhardtii* 80S ribosome have been deposited in the EM and protein database under accession codes EMD-___ (80S ribosome), EMD-____ (60S ribosome), EMD-____ (40S ribosome) and PDB _____ (80S ribosome). Coordinates for previously published structures with newly modeled metal ions are available in the supplementary materials along with associated FEM maps, where applicable.

## Supplementary Materials

Materials and Methods

Figs. S1 to S6

Tables S1 to S2

References (*68–79*)

Movie S1

## References and Notes

1. H. F. Noller, V. Hoffarth, L. Zimniak, Unusual resistance of peptidyl transferase to protein extraction procedures. Science 256, 1416–1419 (1992).

2. N. Ban, P. Nissen, J. Hansen, P. B. Moore, T. A. Steitz, The complete atomic structure of the large ribosomal subunit at 2.4 A resolution. Science 289, 905–920 (2000).

3. A. Korostelev, S. Trakhanov, M. Laurberg, H. F. Noller, Crystal structure of a 70S ribosome-tRNA complex reveals functional interactions and rearrangements. Cell 126, 1065–1077 (2006).

4. M. M. Yusupov et al., Crystal structure of the ribosome at 5.5 A resolution. Science 292, 883–896 (2001).

5. M. Selmer et al., Structure of the 70S ribosome complexed with mRNA and tRNA. Science 313, 1935–1942 (2006).

6. B. T. Wimberly et al., Structure of the 30S ribosomal subunit. Nature 407, 327–339 (2000).

7. A. Sievers, M. Beringer, M. V. Rodnina, R. Wolfenden, The ribosome as an entropy trap. Proc Natl Acad Sci U S A 101, 7897–7901 (2004).

8. P. Nissen, J. Hansen, N. Ban, P. B. Moore, T. A. Steitz, The structural basis of ribosome activity in peptide bond synthesis. Science 289, 920–930 (2000).

9. V. I. Katunin, G. W. Muth, S. A. Strobel, W. Wintermeyer, M. V. Rodnina, Important contribution to catalysis of peptide bond formation by a single ionizing group within the ribosome. Mol Cell 10, 339–346 (2002).

10. G. W. Muth, L. Ortoleva-Donnelly, S. A. Strobel, A single adenosine with a neutral pKa in the ribosomal peptidyl transferase center. Science 289, 947–950 (2000).

11. G. Wallin, J. Aqvist, The transition state for peptide bond formation reveals the ribosome as a water trap. Proc Natl Acad Sci U S A 107, 1888–1893 (2010).

12. Y. S. Polikanov, T. A. Steitz, C. A. Innis, A proton wire to couple aminoacyl-tRNA accommodation and peptide-bond formation on the ribosome. Nat Struct Mol Biol 21, 787–793 (2014).

13. D. M. Lilley, in Philos Trans R Soc Lond B Biol Sci. (England, 2011), vol. 366, pp. 2910–2917.

14. P. L. Adams, M. R. Stahley, A. B. Kosek, J. Wang, S. A. Strobel, Crystal structure of a self-splicing group I intron with both exons. Nature 430, 45–50 (2004).

15. M. R. Stahley, S. A. Strobel, Structural evidence for a two-metal-ion mechanism of group I intron splicing. Science 309, 1587–1590 (2005).

16. A. R. Robart, R. T. Chan, J. K. Peters, K. R. Rajashankar, N. Toor, Crystal structure of a eukaryotic group II intron lariat. Nature 514, 193–197 (2014).

17. N. Toor, K. S. Keating, S. D. Taylor, A. M. Pyle, Crystal structure of a self-spliced group II intron. Science 320, 77–82 (2008).

18. A. M. Lambowitz, S. Zimmerly, Group II introns: mobile ribozymes that invade DNA. Cold Spring Harb Perspect Biol 3, a003616 (2011).

19. A. E. Hesslein et al., Exploration of the conserved A+C wobble pair within the ribosomal peptidyl transferase center using affinity purified mutant ribosomes. Nucleic Acids Res 32, 3760–3770 (2004).

20. K. Lang, M. Erlacher, D. N. Wilson, R. Micura, N. Polacek, The role of 23S ribosomal RNA residue A2451 in peptide bond synthesis revealed by atomic mutagenesis. Chem Biol 15, 485–492 (2008).

21. M. M. Harding, Geometry of metal-ligand interactions in proteins. Acta Crystallogr D Biol Crystallogr 57, 401–411 (2001).

22. H. Zheng, I. G. Shabalin, K. B. Handing, J. M. Bujnicki, W. Minor, Magnesium-binding architectures in RNA crystal structures: validation, binding preferences, classification and motif detection. Nucleic Acids Res 43, 3789–3801 (2015).

23. M. Egli, S. Portmann, N. Usman, RNA hydration: a detailed look. Biochemistry 35, 8489–8494 (1996).

24. E. N. Baker, R. E. Hubbard, Hydrogen bonding in globular proteins. Prog Biophys Mol Biol 44, 97–179 (1984).

25. P. V. Afonine et al., FEM: feature-enhanced map. Acta Crystallogr D Biol Crystallogr 71, 646–666 (2015).

26. P. D. Adams et al., PHENIX: a comprehensive Python-based system for macromolecular structure solution. Acta Crystallogr D Biol Crystallogr 66, 213–221 (2010).

27. J. Wang, Z. Liu, J. Frank, P. B. Moore, Identification of ions in experimental electrostatic potential maps. IUCrJ 5, 375–381 (2018).

28. J. Wang, S. K. Natchiar, P. B. Moore, B. P. Klaholz, Identification of Mg^2+^ ions next to nucleotides in cryo-EM maps using electrostatic potential maps. Acta Crystallogr D Struct Biol 77, 534–539 (2021).

29. E. Sehgal et al., High-resolution reconstruction of a C. elegans ribosome sheds light on evolutionary dynamics and tissue specificity. RNA 30, 1513–1528 (2024).

30. J. Smirnova et al., Structure of the actively translating plant 80S ribosome at 2.2 Å resolution. Nat Plants 9, 987–1000 (2023).

31. P. Tesina et al., Molecular mechanism of translational stalling by inhibitory codon combinations and poly(A) tracts. EMBO J 39, e103365 (2020).

32. P. R. Bhatt et al., Structural basis of ribosomal frameshifting during translation of the SARS-CoV-2 RNA genome. Science 372, 1306–1313 (2021).

33. S. A. Fromm et al., The translating bacterial ribosome at 1.55 Å resolution generated by cryo-EM imaging services. Nat Commun 14, 1095 (2023).

34. S. Trobro, J. Aqvist, Mechanism of peptide bond synthesis on the ribosome. Proc Natl Acad Sci U S A 102, 12395–12400 (2005).

35. J. L. Hansen, T. M. Schmeing, P. B. Moore, T. A. Steitz, Structural insights into peptide bond formation. Proc Natl Acad Sci U S A 99, 11670–11675 (2002).

36. T. M. Schmeing, K. S. Huang, D. E. Kitchen, S. A. Strobel, T. A. Steitz, Structural insights into the roles of water and the 2’ hydroxyl of the P site tRNA in the peptidyl transferase reaction. Mol Cell 20, 437–448 (2005).

37. T. A. Steitz, J. A. Steitz, A general two-metal-ion mechanism for catalytic RNA. Proc Natl Acad Sci U S A 90, 6498–6502 (1993).

38. M. Nowotny, W. Yang, Stepwise analyses of metal ions in RNase H catalysis from substrate destabilization to product release. EMBO J 25, 1924–1933 (2006).

39. J. Hang, R. Wan, C. Yan, Y. Shi, Structural basis of pre-mRNA splicing. Science 349, 1191–1198 (2015).

40. W. P. Galej et al., Cryo-EM structure of the spliceosome immediately after branching. Nature 537, 197–201 (2016).

41. K. Bertram et al., Cryo-EM structure of a human spliceosome activated for step 2 of splicing. Nature 542, 318–323 (2017).

42. R. T. Chan et al., Structural basis for the second step of group II intron splicing. Nat Commun 9, 4676 (2018).

43. K. Eysmont, K. Matylla-Kulińska, A. Jaskulska, M. Magnus, M. M. Konarska, Rearrangements within the U6 snRNA Core during the Transition between the Two Catalytic Steps of Splicing. Mol Cell 75, 538–548.e533 (2019).

44. H. Pelletier, M. R. Sawaya, A. Kumar, S. H. Wilson, J. Kraut, Structures of ternary complexes of rat DNA polymerase beta, a DNA template-primer, and ddCTP. Science 264, 1891–1903 (1994).

45. D. Wang, D. A. Bushnell, K. D. Westover, C. D. Kaplan, R. D. Kornberg, Structural basis of transcription: role of the trigger loop in substrate specificity and catalysis. Cell 127, 941–954 (2006).

46. Y. W. Yin, T. A. Steitz, The structural mechanism of translocation and helicase activity in T7 RNA polymerase. Cell 116, 393–404 (2004).

47. D. A. Hiller, V. Singh, M. Zhong, S. A. Strobel, A two-step chemical mechanism for ribosome-catalysed peptide bond formation. Nature 476, 236–239 (2011).

48. J. S. Weinger, K. M. Parnell, S. Dorner, R. Green, S. A. Strobel, Substrate-assisted catalysis of peptide bond formation by the ribosome. Nat Struct Mol Biol 11, 1101–1106 (2004).

49. S. Dorner, N. Polacek, U. Schulmeister, C. Panuschka, A. Barta, Molecular aspects of the ribosomal peptidyl transferase. Biochem Soc Trans 30, 1131–1136 (2002).

50. H. S. Zaher, J. J. Shaw, S. A. Strobel, R. Green, The 2’-OH group of the peptidyl-tRNA stabilizes an active conformation of the ribosomal PTC. EMBO J 30, 2445–2453 (2011).

51. S. Dorner, C. Panuschka, W. Schmid, A. Barta, Mononucleotide derivatives as ribosomal P-site substrates reveal an important contribution of the 2’-OH to activity. Nucleic Acids Res 31, 6536–6542 (2003).

52. K. S. Huang, N. Carrasco, E. Pfund, S. A. Strobel, Transition state chirality and role of the vicinal hydroxyl in the ribosomal peptidyl transferase reaction. Biochemistry 47, 8822–8827 (2008).

53. T. M. Schmeing, K. S. Huang, S. A. Strobel, T. A. Steitz, An induced-fit mechanism to promote peptide bond formation and exclude hydrolysis of peptidyl-tRNA. Nature 438, 520–524 (2005).

54. M. D. Erlacher et al., Chemical engineering of the peptidyl transferase center reveals an important role of the 2’-hydroxyl group of A2451. Nucleic Acids Res 33, 1618–1627 (2005).

55. M. D. Erlacher et al., Efficient ribosomal peptidyl transfer critically relies on the presence of the ribose 2’-OH at A2451 of 23S rRNA. J Am Chem Soc 128, 4453–4459 (2006).

56. M. Amort et al., An intact ribose moiety at A2602 of 23S rRNA is key to trigger peptidyl-tRNA hydrolysis during translation termination. Nucleic Acids Res 35, 5130–5140 (2007).

57. A. Chirkova et al., The role of the universally conserved A2450-C2063 base pair in the ribosomal peptidyl transferase center. Nucleic Acids Res 38, 4844–4855 (2010).

58. J. A. Piccirilli, T. S. McConnell, A. J. Zaug, H. F. Noller, T. R. Cech, Aminoacyl esterase activity of the Tetrahymena ribozyme. Science 256, 1420–1424 (1992).

59. B. Zhang, T. R. Cech, Peptide bond formation by in vitro selected ribozymes. Nature 390, 96–100 (1997).

60. B. Zhang, T. R. Cech, Peptidyl-transferase ribozymes: trans reactions, structural characterization and ribosomal RNA-like features. Chem Biol 5, 539–553 (1998).

61. R. S. Jansen et al., N-lactoyl-amino acids are ubiquitous metabolites that originate from CNDP2-mediated reverse proteolysis of lactate and amino acids. Proc Natl Acad Sci U S A 112, 6601–6606 (2015).

62. M. D. Moya-Garzon et al., A β-hydroxybutyrate shunt pathway generates anti-obesity ketone metabolites. Cell 188, 175–186.e120 (2025).

63. J. L. Hougland, A. V. Kravchuk, D. Herschlag, J. A. Piccirilli, Functional identification of catalytic metal ion binding sites within RNA. PLoS Biol 3, e277 (2005).

64. M. Boudvillain, A. M. Pyle, Defining functional groups, core structural features and inter-domain tertiary contacts essential for group II intron self-splicing: a NAIM analysis. EMBO J 17, 7091–7104 (1998).

65. G. Chanfreau, A. Jacquier, Catalytic site components common to both splicing steps of a group II intron. Science 266, 1383–1387 (1994).

66. P. M. Gordon, J. A. Piccirilli, Metal ion coordination by the AGC triad in domain 5 contributes to group II intron catalysis. Nat Struct Biol 8, 893–898 (2001).

67. S. M. Fica et al., RNA catalyses nuclear pre-mRNA splicing. Nature 503, 229–234 (2013).

68. N. Sueoka, Mitotic replication of deoxyribonucleic acid in Chlamydomonas reinhardi. Proc Natl Acad Sci U S A 46, 83–91 (1960).

69. S. H. Scheres, RELION: implementation of a Bayesian approach to cryo-EM structure determination. J Struct Biol 180, 519–530 (2012).

70. D. Kimanius, B. O. Forsberg, S. H. Scheres, E. Lindahl, Accelerated cryo-EM structure determination with parallelisation using GPUs in RELION-2. Elife 5, (2016).

71. S. Q. Zheng et al., MotionCor2: anisotropic correction of beam-induced motion for improved cryo-electron microscopy. Nat Methods 14, 331–332 (2017).

72. A. Punjani, J. L. Rubinstein, D. J. Fleet, M. A. Brubaker, cryoSPARC: algorithms for rapid unsupervised cryo-EM structure determination. Nat Methods 14, 290–296 (2017).

73. S. F. Altschul, W. Gish, W. Miller, E. W. Myers, D. J. Lipman, Basic local alignment search tool. J Mol Biol 215, 403–410 (1990).

74. E. F. Pettersen et al., UCSF Chimera--a visualization system for exploratory research and analysis. J Comput Chem 25, 1605–1612 (2004).

75. P. Emsley, K. Cowtan, Coot: model-building tools for molecular graphics. Acta Crystallogr D Biol Crystallogr 60, 2126–2132 (2004).

76. K. S. Keating, A. M. Pyle, Semiautomated model building for RNA crystallography using a directed rotameric approach. Proc Natl Acad Sci U S A 107, 8177–8182 (2010).

77. K. S. Keating, A. M. Pyle, RCrane: semi-automated RNA model building. Acta Crystallogr D Biol Crystallogr 68, 985–995 (2012).

78. P. V. Afonine et al., Real-space refinement in PHENIX for cryo-EM and crystallography. Acta Crystallogr D Struct Biol 74, 531–544 (2018).

79. A. Morin et al., Collaboration gets the most out of software. Elife 2, e01456 (2013).

