## Supplementary Information for "Structural evidence for metal ion catalysis in the ribosome"

**The PDF file includes:**

Materials and Methods
Figs. S1 to S6
Tables S1 to S2
References

**Other Supplementary Materials for this manuscript include the following:**

Movie S1

### Materials and Methods

#### *C. reinhardtii* ribosome purification

A 3-L culture of *C. reinhardtii* strain CC-400 was grown autotrophically in Sueoka's HS media (*68*) at 22 °C with 200 rpm orbital shaking, to a density of 1.8 x $10^7$ cells/mL as determined by hemocytometer count. All subsequent steps were carried out at 4 °C. Cells were pelleted by centrifugation at 5,000 rcf for 10 minutes. The cell pellet was resuspended in 50 mL lysis buffer (150 mM NaCl, 60 mM KCl, 10 mM $MgCl_2$, 0.0085% n-dodecyl-β-D-maltoside, 100 mM HEPES, pH 8.0) with Protease Inhibitor Cocktail VI (Millipore) added immediately prior to use. Cells were lysed using a probe sonicator for 2 minutes of total run time at 40% amplitude, using 10-second bursts and 60-second pauses, and cell debris was cleared by centrifuging at 2,000 rcf for 30 min. The resulting supernatant was further cleared by ultracentrifugation at 40,000 rcf for 45 min. Bis(sulfosuccinimidyl) suberate was then added to a working concentration of 1 mM in a 1-mL aliquot of the supernatant. The sample was loaded onto a 12-mL 10–50% sucrose gradient in lysis buffer, and ultracentrifuged at 115,000 rcf for 17 hours. The gradient was then fractionated into 1-mL aliquots, and 50 µL from the bottommost fraction of the gradient was passed through a mini Quick Spin RNA Column (Roche).

#### Single-particle cryo-EM sample preparation

3.5 µL of the purified ribosome sample was loaded onto a Quantifoil Cu300 R2/2 + 2 nm C film grid that had been glow discharged using a PELCO easiGlow (0.4 mbar, 15 mA for 30 sec). The grid was blotted for 5 sec with filter paper (Whatman No.1) and then immediately plunged into a liquid ethane/propane (37.5/62.5) mix using a manual plunger.

#### Single particle cryo-EM data collection and processing

4032 movies were collected on a Titan Krios microscope (Thermo Fisher) operating at 300 keV, equipped with a Gatan K3 direct electron detector, and with a magnification of 81,000x for a physical pixel size of 1.05 Angstroms and camera operating in super-resolution mode (0.524 Å/pixel). Movies were collected as 50 dose-fractionated frames for 7.3 s with a dose rate of 7.5 $e^-$/px/s, for a total dose of 50 $e^-/Å^2$, with a nominal defocus range of -1.0 to -2.5 microns.

Super resolution movies were imported into RELION 3.1 (*69, 70*) and motion corrected with a pixel binning of 2 to make the pixel size 1.05 Å/px, using RELION's internal version of MotionCor2 (*71*). These 4032 exposures were then imported into cryoSPARC 4.1.2 (*72*), and CTF-corrected using cryoSPARC's Patch CTF. 780,255 particles were picked on 4,032 micrographs using blob picker with a minimum particle diameter of 250 Å and a maximum particle diameter of 325 Å. 130 of these micrographs and corresponding 23,291 particles were manually removed based on poor CTF fit.

The cryo-workflow methods is described in **Fig. S6**. In brief, the 756,964 particles were extracted with a 144 pixel box (3.14 Å/px). Particles close to the edge of the micrograph were automatically discarded, and the remaining 595,520 particles were subjected to a single round of 2D classification. Particles from 2D classes that looked like the ribosome were selected for a 2-class ab initio model generation. The volumes generated here were then used for a single round of heterogenous classification. These particles were used for homogenous refinement, which was used to generate a set of 2D templates for cryoSPARC's template particle picker.

695,548 particles from template picker and the 164,511 particles from blob picker were extracted with a 180 pixel box (3.14 Å/px), and put through several rounds of 2D and heterogenous refinement for 3D classification. The remaining 345,502 particles were put through refinement, and re-extracted with a 432 pixel box (1.31 Å/px). In order to remove duplicate particles from the combined blob and template picking, particles with 3D alignment centers within 50 Å were removed. These particles were then extracted with an unbinned 540 pixel box (1.05 Å/px), and subjected to another round of duplicate removal. These 201,007 particles were then refined, and classified using cryoSPARC's 3D classification algorithm, using the default size of 10 classes. The class with no E-site tRNA was selected for non-uniform refinement. These 71,404 particles selected were refined to 2.73 Å after non-uniform refinement.

<u>rRNA and protein sequences</u>

Ribosomal RNA and protein sequences were retrieved by running corresponding sequences from the *N. tabacum* ribosome (based on those available in a published structure, PDB ID: 8B2L) through NCBI's BLAST (*73*) and restricting results to the organism *Chlamydomonas reinhardtii*. The sequence for eL39 could not be found via BLAST searches of the NCBI protein database, and was instead identified via a BLAST search of the UniParc archive within the UniProt database (https://www.uniprot.org/).

<u>*C. reinhardtii* ribosome model building</u>

Modeling of the 80S ribosome of *C. reinhardtii* was performed by first using UCSF Chimera (*74*) to manually dock a model of the *S. cerevisiae* and *N. tabacum* 80S ribosomes (PDB IDs 6WOO and 8B2L, respectively) into the *C. reinhardtii* electron density map, followed by use of the "fit in map" tool, for ease of identification of rRNA sequence register and ribosomal protein locations. Modeling of rRNA was then performed in Coot (*75*) using the RCrane (*76, 77*) plugin by removing the docked rRNA content after identifying the sequence register and then modeling into the empty density. The 3′CCA of the P-site tRNA was modeled into the density *de novo*. Ribosomal proteins were modeled in Coot by mutating the docked proteins and using real space refinement of small stretches (~5–20 amino acids) to precisely place each residue. The 60S and 40S subunits were modeled from two distinct maps that exhibited slightly higher resolution for the corresponding subunit compared to the 80S map, and additional local refinement maps produced in cryoSPARC were used to facilitate modeling at various regions around the periphery of the ribosome. Upon completion of model building, the 60S and 40S models were each fit into the density of the 80S map using the "fit in map" tool in Chimera to generate a single 80S coordinate file.

<u>Structure refinement and statistical analysis</u>

Structure refinement and calculation of feature-enhanced maps were performed using Phenix (*26, 78*). To calculate per-nucleotide all-atom RMSD values, models of the triplex from each structure indicated were superposed with the triplex from *C. reinhardtii* in Coot using LSQ superpose, and the superposed models were then opened in UCSF Chimera for analysis. The active site residues were remodeled in the cryo-EM structures to better fit the density using real-space refinement in Coot. For the 80S *C. reinhardtii* ribosome, per-chain rigid body refinement was performed using Phenix. Real space refinement in Coot was used to address clashes, followed by real space refinement in Phenix. Additional clashes were fixed via real space refinement in Coot, followed by a final round of rigid body refinement in Phenix. All map/model validation and statistics were done in Phenix (*26, 78*) (**Table S2**). All software was compiled by SBGrid (*79*).

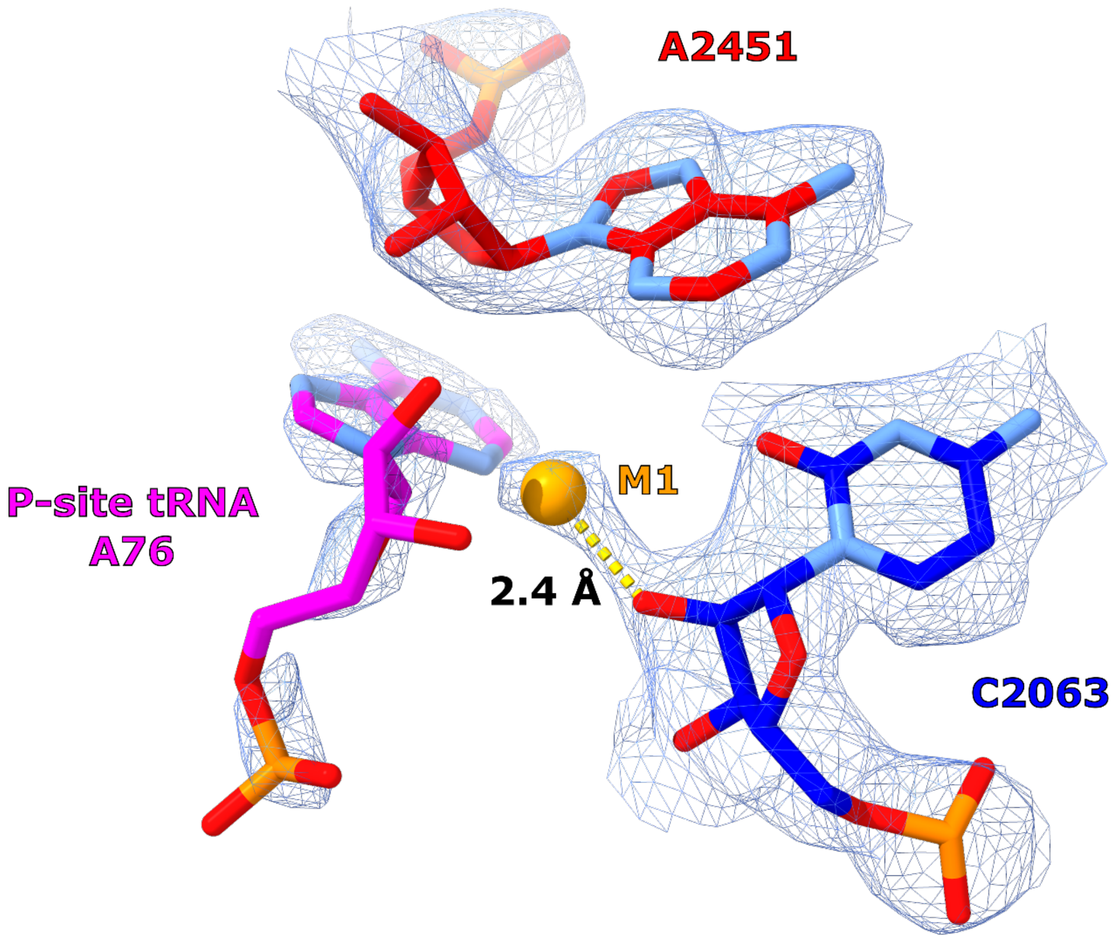

**Fig. S1. Local architecture of M1 binding site.** Cryo-EM structure from *C. reinhardtii* showing M1 (orange) coordinating with the 2′-OH of C2063 (blue) from the large subunit rRNA. A2451 (red) from the large subunit rRNA and A76 of the P-site tRNA (magenta) are in the immediate vicinity of the metal ion.

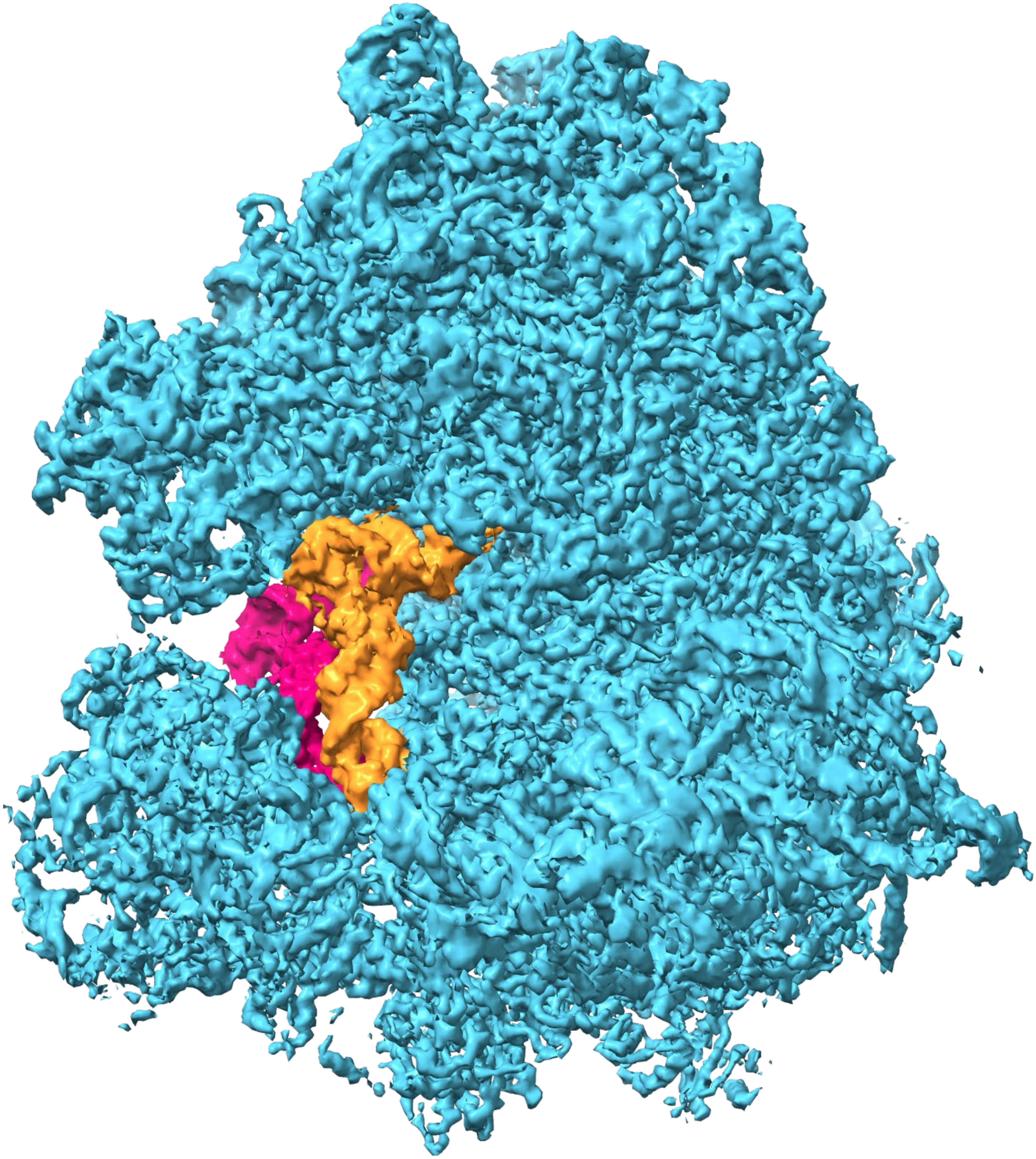

**Fig. S2. Cryo-EM structure of *Chlamydomonas reinhardtii* cytosolic ribosome with tRNAs.** A map of the *Chlamydomonas reinhardtii* ribosome after performing focused classification on the tRNA. Ribosomal RNA and proteins are in cyan, P-site tRNA in orange, and E-site tRNA in magenta.

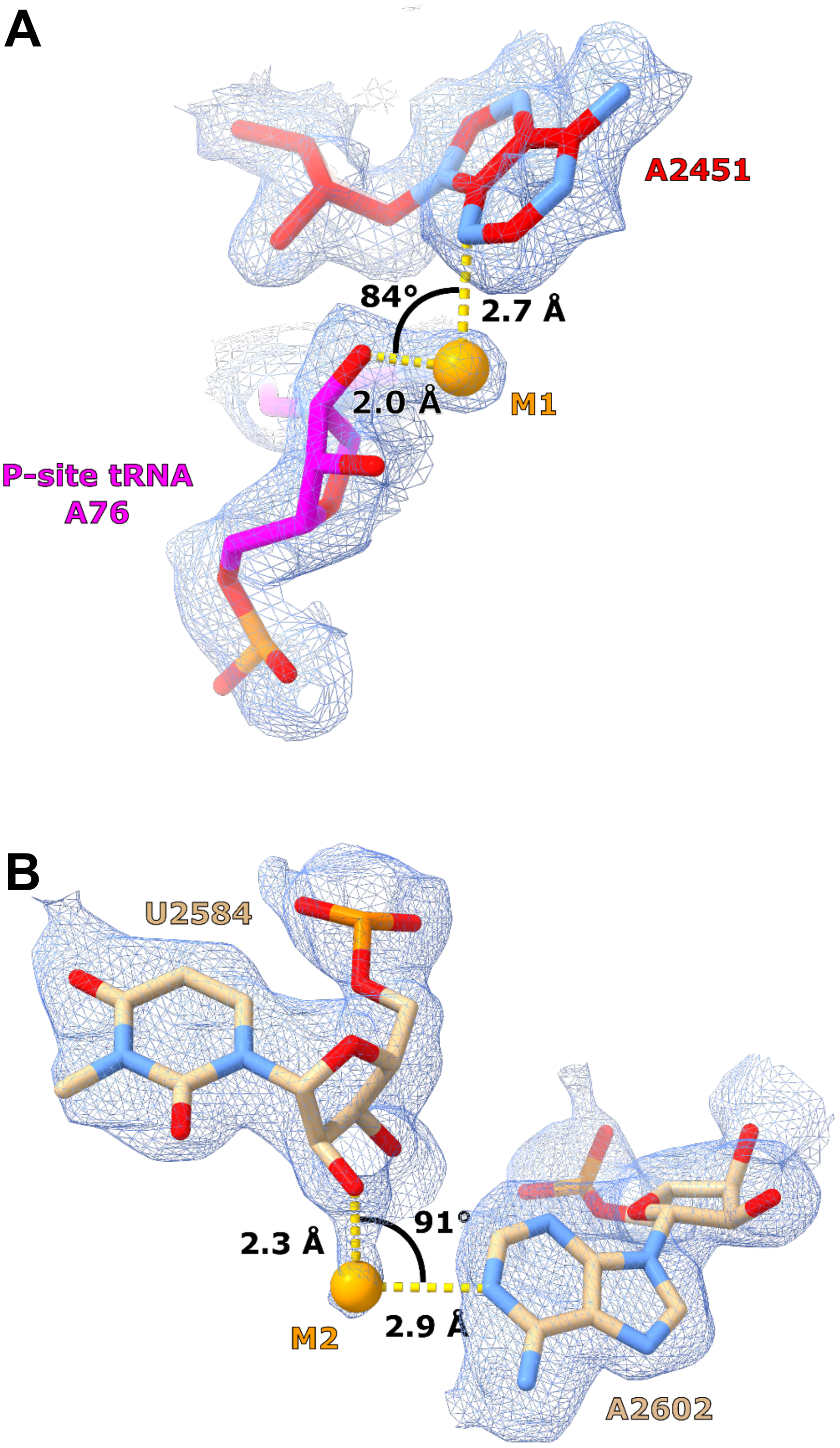

**Fig. S3. Binding geometry of metal ions with respect to 2′-OH and adenosine ligands.** (**A**) When M1 coordinates to the peptidyl-tRNA A76, it is positioned between the 2′-OH of A76 and the N3 of A2451. The angle from the 2′-OH to M1 to the N3 is 84° (PDB 8B0X). (**B**) M2 is positioned between the 2′-OH of U2584 and the N1 of A2602. The angle from the 2′-OH to M2 to the N1 is 91° (PDB 1VQ5, FEM map contoured at 1.5σ).

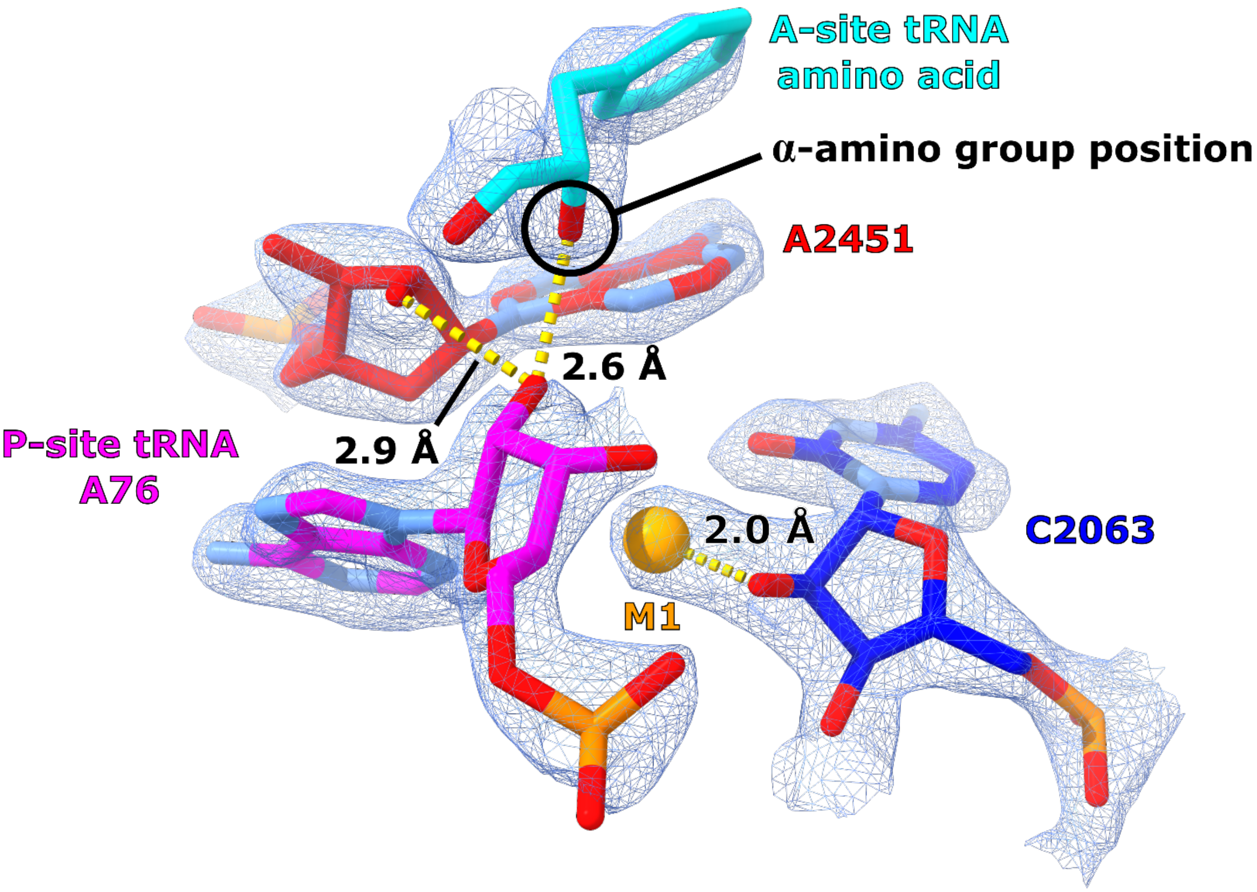

**Fig. S4. The nucleophilic α-amine hydrogen-bonds with the 2′-OH of peptidyl-tRNA A76.** An example from x-ray crystallography (PDB 1VQN, FEM map contoured at 2.3σ) where the atom corresponding to the α-amine nucleophile (a hydroxyl group in this substrate analog) is hydrogen-bonding with the 2′-OH of A76 from the peptidyl-tRNA at a distance of 2.6 Å. The 2′-OH of A2451 is also hydrogen-bonding to the 2′-OH of A76 at a distance of 2.9 Å. M1 is inner-shell coordinated to the 2′-OH of C2063 at a distance of 2.0 Å. Among the crystal structures analyzed in this work, 1VQN is the only one where this view is possible due to a convergence of factors: substrate is bound in the A-site, the ribosome does not contain a tetrahedral intermediate analog, and the P-site A76 is not deoxygenated at the O2′ position.

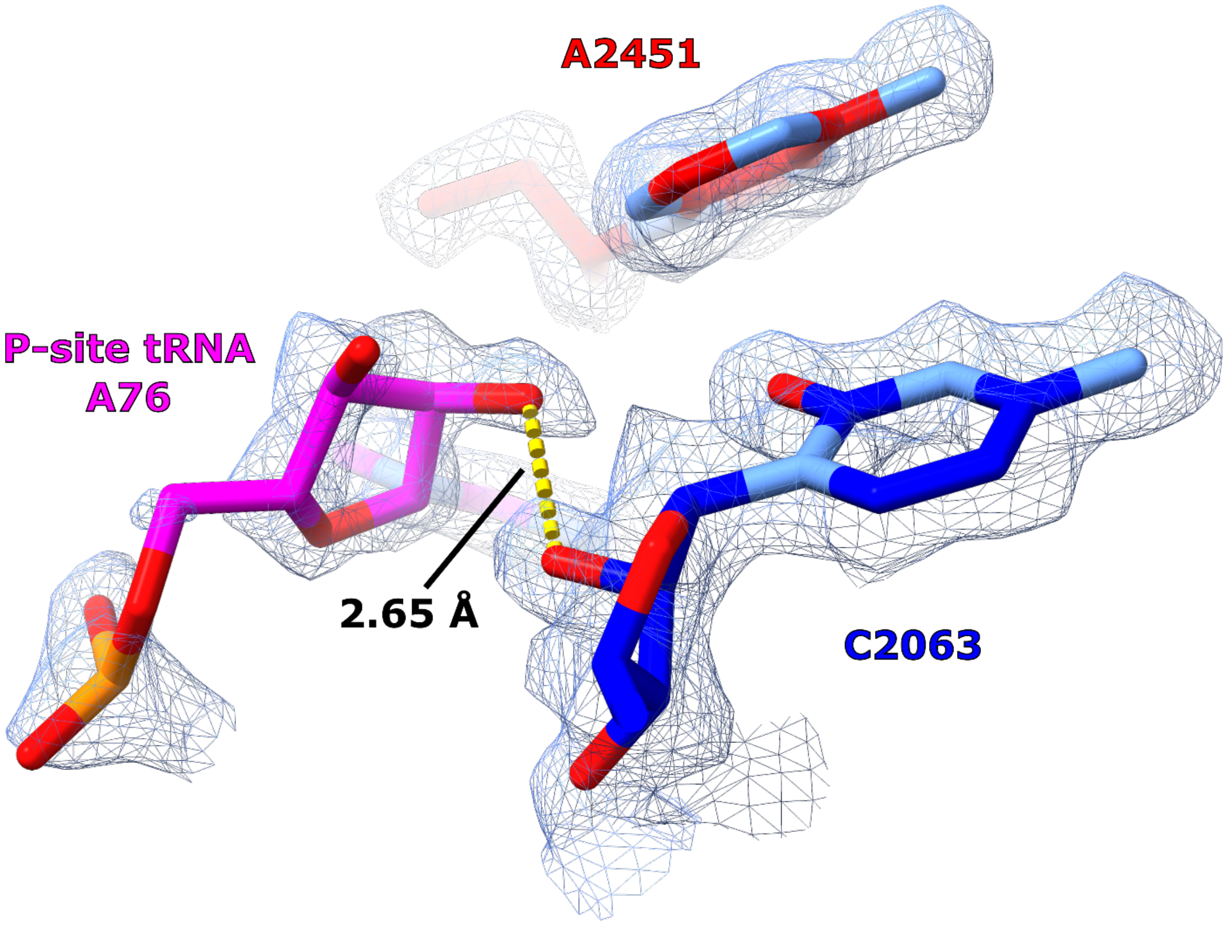

**Fig. S5. The M1 binding site collapses when the 2′-OH of peptidyl-tRNA A76 directly hydrogen-bonds to the 2′-OH of C2063.** In the 1.55 Å map (EMD-15793) published with PDB 8B0X, the electron density indicates a "collapsed" state for the M1 binding site, where the ribose of the P-site A76 is not held in a C2′ endo conformation. The 2′-OH of A76 is not positioned to interact with the 2′-OH of A2451 in this configuration, and is instead hydrogen-bonding with the 2′-OH of C2063, at a distance of 2.65 Å.

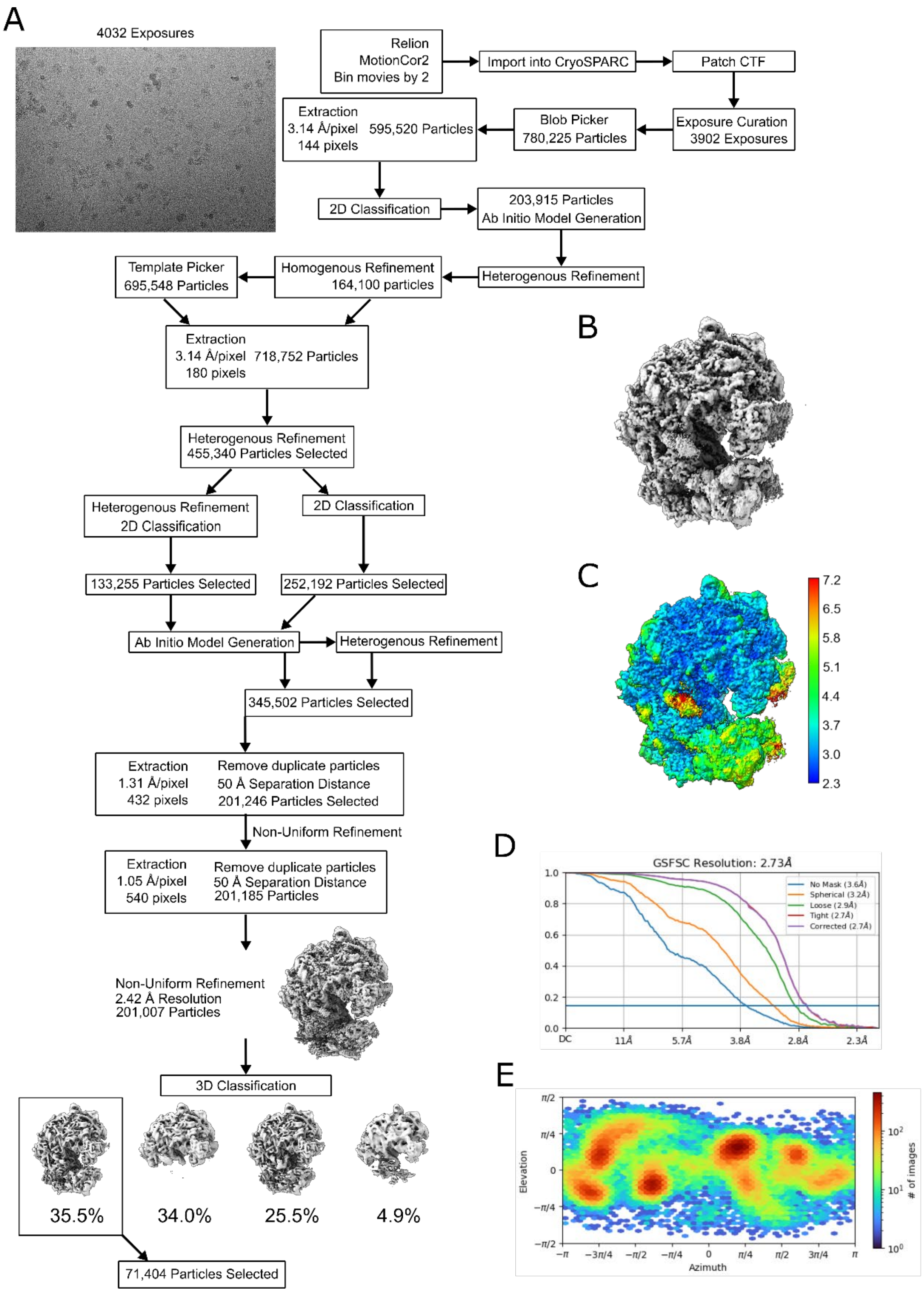

**Fig. S6. Cryo-EM workflow.** (**A**) Cryo-EM workflow for *C. reinhardtii* ribosome. (**B**) Final reconstruction after non-uniform refinement for the selected ribosome class. (**C**) Local resolution map of ribosome. (**D**) GSFSC curve of reconstruction. (**E**) Particle orientation plot for the final reconstruction.

**Table S1. Coordination distances between M1 and the 2′-OH of C2063 or A76.**

| | PDB Accession | 2′-OH distance from M1 (Å) A76 | C2063 |
|---|---|---|---|
| Cryo-EM | 8B0X | **2.02** | 2.85 |
| | 7O7Y | **2.51** | 3.15 |
| | 8B2L | **2.46** | 3.09 |
| | 9BH5 | 4.16 | **2.09** |
| | 7O7Z | 3.21 | **2.54** |
| | *C. reinhardtii* 80S | 3.88 | **2.43** |
| X-ray | 1M90 | 3.01 | **2.21** |
| | 1VQ4 | N/A | **2.13** |
| | 1VQ5 | 3.68 | **2.19** |
| | 1VQL | N/A | **2.23** |
| | 1VQN | 3.64 | **1.98** |

**Table S2. Cryo-EM data collection and refinement statistics.**

| | *C. reinhardtii* 80S ribosome |
|---|---|
| **Data collection** | |
| Microscope | FEI Titan Krios |
| Voltage (kV) | 300 |
| Camera | Gatan K3 Summit |
| Magnification | 81,000 |
| Nominal defocus range (μm) | -1.0/-2.5 |
| Exposure time (s) | 7.324 |
| Number of frames | 50 |
| Dose rate ($e^-$/Pixel/s) | 7.53 |
| Total dose ($e^-$/Å$^2$) | 50 |
| Pixel size (Å) | 1.048 |
| Micrographs collected | 4,032 |
| Micrographs processed | 3,902 |
| Total particles | 201,007 |
| Particles used in final map | 71,404 |
| Map resolution global (FSC 0.143) masked | 2.73 |
| **Model composition** | |
| Non-hydrogen atoms | 194673 |
| Protein residues | 10938 |
| RNA bases | 5028 |
| Ligands | 3 |
| **Refinement (PHENIX)** | |
| Refinement package | **Real space refinement** |
| CC (volume) | 0.85 |
| CC (mask) | 0.87 |
| CC (peak) | 0.76 |
| FCS map-to-model (0.5) masked | 3.01 |
| Map sharpening B-factor (Å$^2$) | -68.8 |
| **Rms deviations** | |
| Bond length (Å) | 0.004 |
| Bond angles (°) | 0.824 |
| **Validation** | |
| Molprobity score | 2.09 |
| All-atom clashscore | 12.14 |

**Movie S1. Catalytic triplex overlay spin movie.**
